# Dysregulation of the basal ganglia indirect pathway prior to cell loss in the Q175 mouse model of Huntington’s disease

**DOI:** 10.1101/2021.01.06.425589

**Authors:** Joshua W. Callahan, David L. Wokosin, Mark D. Bevan

## Abstract

The psychomotor symptoms of Huntington’s disease (HD) are linked to degeneration of the basal ganglia indirect pathway. To determine how this pathway is perturbed prior to cell loss, optogenetic- and reporter-guided electrophysiological interrogation approaches were applied to early symptomatic 6-month-old Q175 HD mice. Although cortical activity was unaffected, indirect pathway striatal projection neurons were hypoactive *in vivo*, consistent with reduced cortical input strength and dendritic excitability. Downstream parvalbumin-expressing prototypic external globus pallidus (GPe) neurons were hyperactive *in vivo* and exhibited elevated autonomous firing *ex vivo*. Optogenetic inhibition of prototypic GPe neurons ameliorated the abnormal hypoactivity of postsynaptic subthalamic nucleus (STN) and putative arkypallidal neurons *in vivo*. In contrast to STN neurons, autonomous arkypallidal activity was unimpaired *ex vivo*. Together with previous studies, these findings demonstrate that basal ganglia indirect pathway neurons are highly dysregulated in Q175 mice through changes in presynaptic activity and/or intrinsic properties 6-12 months before cell loss.

## Introduction

Huntington’s disease (HD) is an autosomal dominant neurodegenerative disorder caused by an expansion of greater than 35 trinucleotide CAG repeats in exon 1 of the huntingtin gene (HTT) (Bates et al., 2015). The age at which HD manifests clinically is inversely related to expansion length, although genetic variability, in for example, capacity for DNA repair, may cause onset to vary by several years in individuals with identical expansions (Andrew et al., 1993; Brinkman et al., 1997; Moss et al., 2017). The cardinal features of HD are the progressive dysfunction and degeneration of the basal ganglia and cortex, which leads to the emergence of debilitating motor, cognitive, and psychiatric symptoms (Vonsattel and DiFiglia, 1998; Bates et al., 2015). The products of the mutant huntingtin gene (mHTT), including both full-length mutant huntingtin (mhtt) and its N-terminal fragment, impair multiple processes that are critical for neuronal function, including axonal transport, autophagy, gene transcription, homeostatic regulation of extracellular glutamate and K^+^, mitochondrial physiology, mTOR signaling, myelination, and synaptic transmission (DiFiglia et al., 1997; Sapp et al., 1999; Gunawardena et al., 2003; Szebenyi et al., 2003; Milnerwood and Raymond, 2010; Reddy and Shirendeb, 2012; Seredenina and Luthi-Carter, 2012; Johri et al., 2013; Pryor et al., 2014; Tong et al., 2014; Lee et al., 2015; Martin et al., 2015; Jiang et al., 2016; Rosas et al., 2018). However, the mechanisms that cause the selective dysregulation, degeneration, and ultimate loss of cortico-basal ganglia circuitry and the relationship of pathogenesis to symptomatology remain poorly understood.

Under normal conditions, the basal ganglia promote the generation of contextually appropriate actions and thoughts, in part through their processing of functionally diverse cortical inputs (Mink and Thach, 1993; Maurice et al., 1999; Tachibana et al., 2008; Cui et al., 2013; Keeler et al., 2014; Nelson and Kreitzer, 2014; Tecuapetla et al., 2016; Markowitz et al., 2018; Klaus et al., 2019). Cortical information is first processed by complex microcircuits within the basal ganglia, including the so-called direct and indirect pathways; the results of these computations are then broadcast to the thalamus, midbrain, and brainstem, affecting behavior (Albin et al., 1989; Gerfen et al., 1990). In adult-onset HD (aoHD), D2 dopamine receptor expressing striatal projection neurons (D2-SPNs) that comprise the first leg of the indirect pathway are relatively susceptible (Albin et al., 1992; Richfield et al., 1995; Sapp et al., 1995). Indeed, early loss of indirect pathway function has been posited to underlie symptoms that manifest in the initial clinical phase of aoHD, including hyperkinesia and loss of behavioral control (Reiner et al., 1988; Vonsattel and DiFiglia, 1998). With disease progression, degeneration of direct pathway D1 dopamine receptor expressing striatal projection neurons (D1-SPNs), together with more widespread degeneration of the cortico-basal ganglia-thalamo-cortical circuit, may underlie bradykinesia and rigidity, and cognitive and psychiatric symptoms that present in mid- to late-stage aoHD (Albin et al., 1990; Deng et al., 2004). In juvenile-onset HD (joHD), which is associated with repeat expansions in excess of 54, bradykinesia, rigidity, and severe psychiatric and cognitive symptoms present in the absence of an early hyperkinetic phase (Fusilli et al., 2018; Tereshchenko et al., 2019), possibly reflecting the relatively widespread susceptibility of cortico-basal ganglia circuitry to more toxic mhtt species. Prior to clinically manifest HD, longitudinal studies also suggest that there is a long, multi-year prodromal period in which subtle but detectible motor, cognitive, and psychiatric deficits are present (Paulsen, 2010; Long et al., 2014; Reilmann et al., 2014).

To better understand HD pathogenesis and pathophysiology, transgenic and knock-in (KI) animal models of HD were developed. The most commonly studied transgenic, N-terminal fragment and full-length mHTT, and KI models possess long repeat sizes that would give rise to joHD in humans (Menalled and Chesselet, 2002; Farshim and Bates, 2018). Consistent with joHD, these mouse models typically exhibit bradykinesia rather than chorea or hyperkinesia (Pouladi et al., 2013; Kosior and Leavitt, 2018). Although these models exhibit inclusion pathology, brain volume loss, and cellular physiological and cognitive deficits analogous to those seen in both aoHD and joHD, frank cell loss is relatively modest (Menalled and Chesselet, 2002; Milnerwood and Raymond, 2010; Raymond et al., 2011). Therefore, it remains unclear whether these models mimic prodromal HD and/or the early stages of clinically manifest aoHD or joHD.

*Ex vivo* electrophysiological analyses of HD models at pre-symptomatic and symptomatic ages have revealed that neurons in the cortico-basal ganglia-thalamo-cortical circuit exhibit complex alterations in their cellular physiological and synaptic properties (Cepeda et al., 2007; Raymond et al., 2011; Plotkin and Surmeier, 2015). In some cases, these changes are the direct result of mhtt in the affected cell, i.e., they are cell autonomous in nature. In other cases, alterations may be triggered by dysregulation of the circuit in which neurons are embedded and are therefore homeostatic or compensatory. The properties of D2-SPNs in HD mice have been studied extensively. Thus, D2-SPNs exhibit loss of cortico-striatal long-term potentiation (LTP), and reductions in axospinous synapse density, mEPSC amplitude, and dendritic excitability (Plotkin et al., 2014; Plotkin and Surmeier, 2014; Sebastianutto et al., 2017; Carrillo-Reid et al., 2019). LTP loss and dendritic hypoexcitability have been attributed to abnormal upregulation of postsynaptic PTEN signaling (Plotkin et al., 2014; Plotkin and Surmeier, 2014). Furthermore, LTP loss and dendritic hypoexcitability may be largely cell autonomous in nature because they can be reversed by lowering mhtt in D2-SPNs (Carrillo-Reid et al., 2019). These *ex vivo* observations suggest that the indirect pathway will be less effectively engaged by cortical excitation in HD mice. Consistent with this view, the activities of cortical neurons and SPNs *in vivo* are less well correlated in HD mice (Miller et al., 2008; Walker et al., 2008; Estrada-Sanchez et al., 2015). However, downstream external globus pallidus (GPe) neurons exhibit no sign of disinhibition *in vivo*, i.e. they are not hyperactive (Beaumont et al., 2016). Thus, the effects of cortical drive on downstream components of the basal ganglia cannot be easily predicted from *ex vivo* observations alone. Furthermore, the above-mentioned *in vivo* recording studies are difficult to interpret due to the heterogeneous nature of cell types in the recorded nuclei and absence of approaches that could establish the identity of recorded neurons.

Thus, the aim of this study was to determine how mhtt expression affects the cortical patterning of identified neurons in the basal ganglia, specifically focusing on components of the indirect pathway: D2-SPNs, prototypic GPe neurons, and subthalamic nucleus (STN) neurons. To address this aim, we utilized a well-characterized mouse model of HD, the Q175 KI model, which expresses human HTT exon 1 with ∼190 trinucleotide repeats in the endogenous HTT gene, and recapitulates many of the progressive molecular, neuropathological, and behavioral abnormalities seen in HD patients (Heikkinen et al., 2012; Menalled et al., 2012). We then applied electrophysiological recording techniques to compare neuronal activity in wild type (WT) and HD mice *in vivo*. To optogenetically identify and manipulate the activity of specific cortical and basal ganglia cell classes, cre recombinase-dependent, viral-mediated expression of channelrhodopsin (ChR2(H134R)) or archaerhodopsin (Arch) was employed. In pilot experiments, we found that the stereotyped patterns of cortical activity present under urethane anesthesia were similar in WT and Q175 mice, enabling us to compare their impact on basal ganglia activity. In the urethane-anesthetized preparation, the cortex exhibits robust ∼1 Hz slow-wave activity (SWA) during which cortical projection neurons, including those projecting to the basal ganglia, exhibit synchronous transitions between hyperpolarized quiescent and depolarized active states (Magill et al., 2001; Walters et al., 2007; Mallet et al., 2008a; Mallet et al., 2008b; Zold et al., 2012; Sharott et al., 2017; Kovaleski et al., 2020). This stereotyped pattern of cortical activity is analogous to that occurring during deep sleep (Steriade, 2000). In addition, somatosensory stimulation was used to trigger cortical activation (ACT), in order to probe the impact of persistent, desynchronized cortical activity (analogous to that seen during arousal) on downstream basal ganglia activity (Steriade, 2000). Finally, we utilized patch clamp recording *ex vivo* to examine whether the altered activity of GPe neurons in Q175 mice *in vivo* was due in part to alterations in autonomous firing. Together with other studies, our data argue that mhtt profoundly dysregulates cortico-basal ganglia circuit dynamics prior to major cell loss through both cell-autonomous and non-cell autonomous mechanisms. These findings could be informative for the design and interpretation of completed, ongoing, and future clinical trials of mhtt-lowering therapeutics.

## Results

Data are reported as median and interquartile range. Data are represented graphically as violin (kernel density) plots and overlaid box plots, with the median (central line), interquartile range (box), and 10-90% range (whiskers) denoted. To minimize assumptions concerning the distribution of data, non-parametric, two-tailed statistical comparisons were made using the Mann-Whitney U (MWU) and Wilcoxon signed-rank (WSR) tests for unpaired and paired comparisons, respectively. In addition, Fisher’s exact test was used for contingency analyses. P < 0.05 was considered significant. Where appropriate, P values were adjusted for multiple comparisons using the Holm-Bonferroni method. Plots and statistical comparisons were generated in Prism (GraphPad Software, Inc., La Jolla, CA, USA; RRID: SCR_002798) and R (https://www.r-project.org/;RRID:SCR_001905).

### Cortical activity is similar in Q175 and WT mice during both SWA and ACT

To determine whether cortico-basal ganglia circuit activity is dysregulated in 6-month-old Q175 mice relative to WT age-matched controls, neuronal activity was compared under urethane anesthesia during both cortical SWA and sensory-evoked cortical ACT. Cortical network activity was assessed from the intracranial electroencephalogram (EEG), which was obtained from a peridural screw “electrode” affixed over primary motor cortex. Cortical SWA was manifest in the EEG as a high-amplitude, low-frequency (∼1 Hz) oscillation upon which phase-locked, low-amplitude, high-frequency oscillations were superimposed (**Figure 1A**). Cortical ACT occurred spontaneously or could be triggered by somatosensory stimulation, and was signified in the EEG by diminution of the 1 Hz oscillation and persistence of higher frequency oscillations (**Figure 1A**). During cortical SWA, the power of motor cortical oscillations in frequency bands ranging from 0-100 Hz were similar in Q175 and WT mice (**Figure 1B-E; Table 1**). In addition, low-frequency and high-frequency cortical oscillations were attenuated and elevated, respectively, to a similar degree in Q175 and WT mice during hind paw pinch-evoked cortical ACT (**Figure 1B-E; Table 1**). Together, these data suggest that motor cortical network activity is similar in 6-month-old Q175 and WT mice.

**Table 1.**
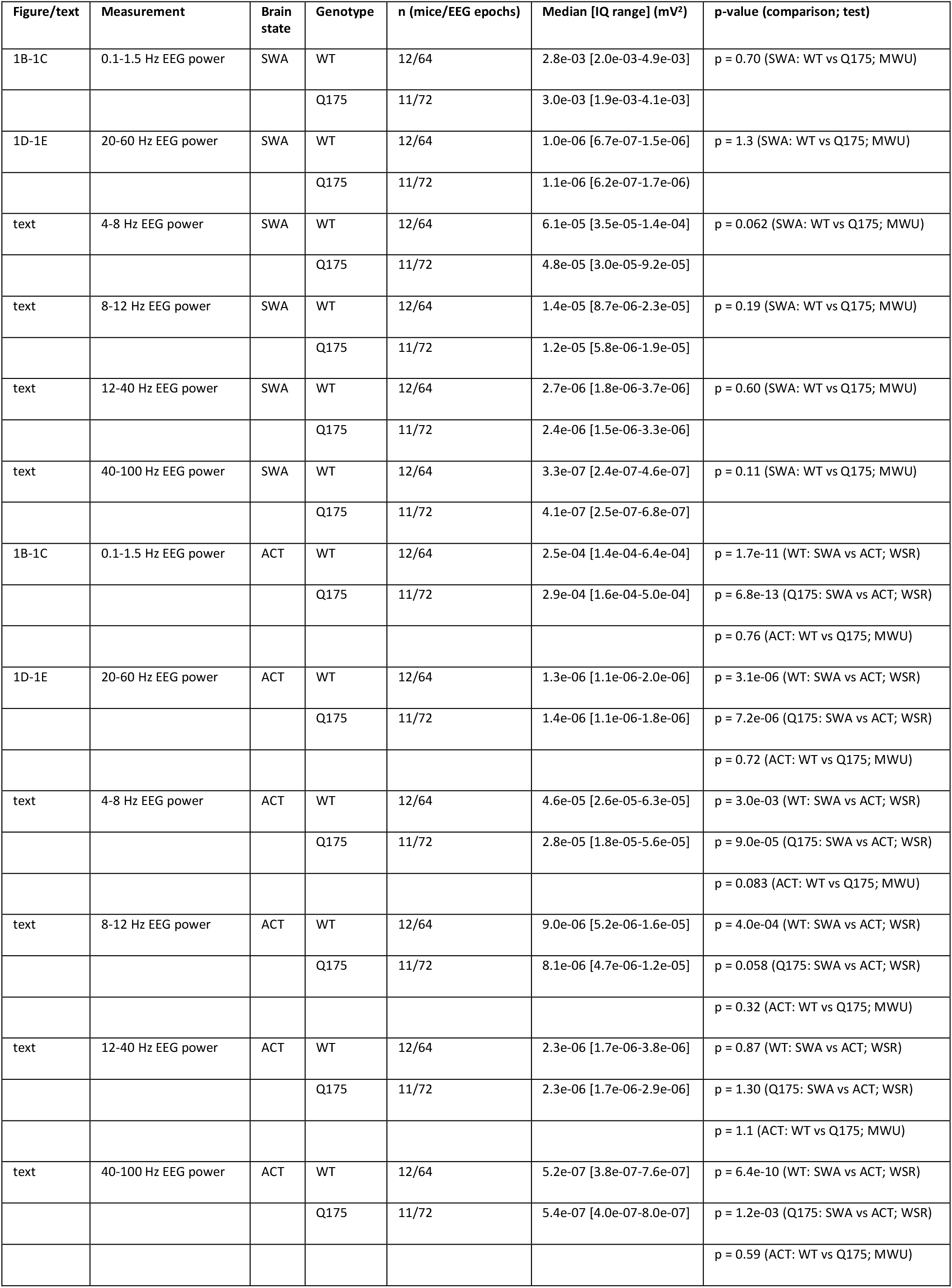

**Figure 1.**
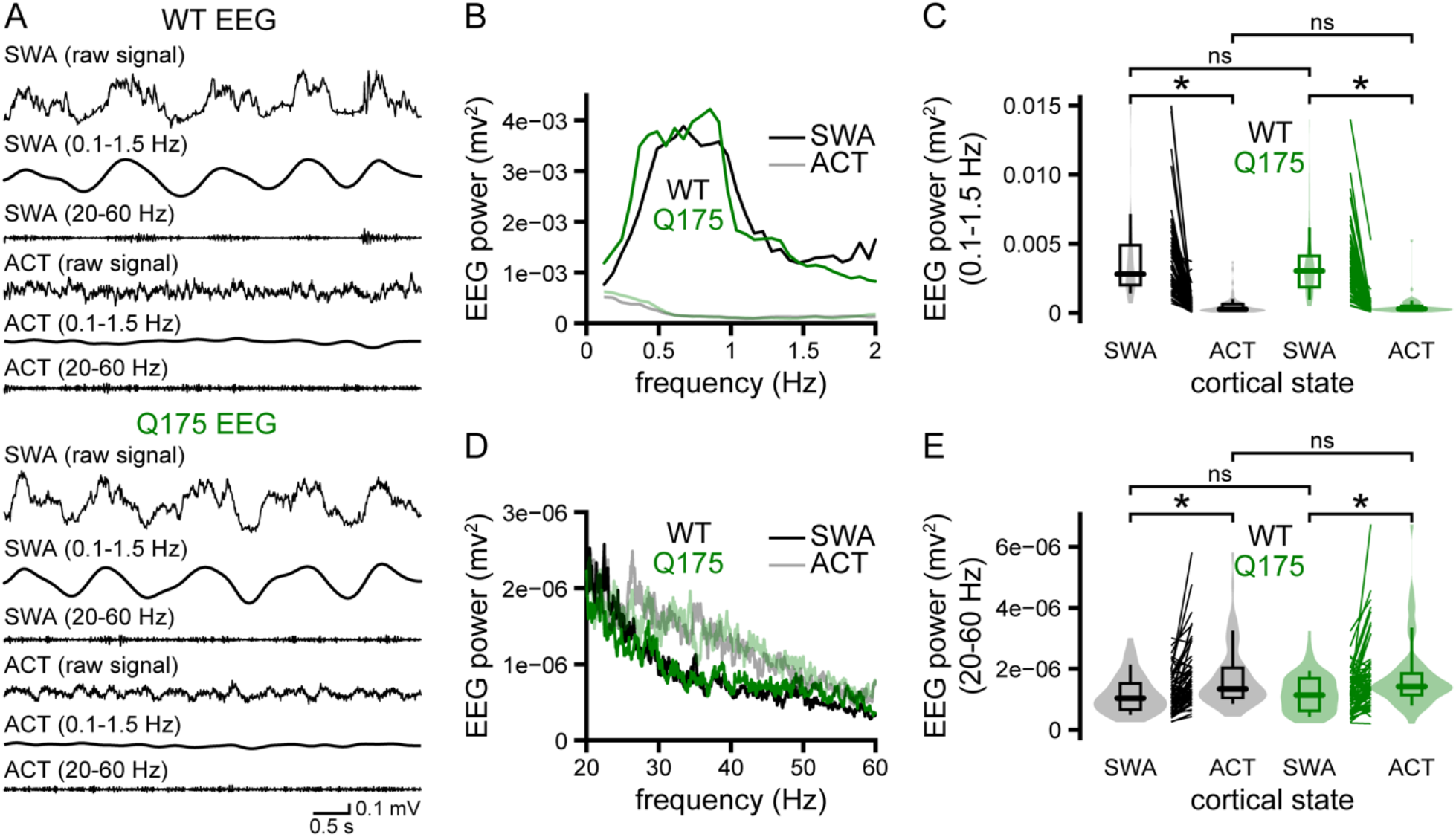
Motor cortical network activity is similar in Q175 and WT mice during both SWA and ACT. (**A-E**) The motor cortical EEGs of urethane-anesthetized Q175 and WT mice were similar during SWA and ACT (**A**, examples; **B-E**, population data; WT, n = 12 mice; Q175, n = 11 mice). Spectral power was similar for both low-(**B, C**) and high-frequency (**D, E**) EEG components. *, p < 0.05. ns, not significant.

Layer V cortical pyramidal neurons, some of which innervate the basal ganglia, are comprised of two major cell classes, pyramidal tract type (PT) and intratelencephalic type (IT) neurons (Harris and Shepherd, 2015). To compare the activity of PT neurons in Q175 and WT mice, an “opto-tagging” approach was employed to identify their firing *in vivo*. First, ChR2(H134R)-eYFP was virally expressed in PT cortical neurons through injection of an adeno-associated virus (AAV) carrying a cre-dependent expression construct into the primary motor cortex of Q175 and WT mice that had been crossed with a PT neuron selective cre-driver line (PT-kj18-cre) (**Figure 2A-C**). 2-4 weeks later, the activity of cortical neurons was compared using an array of tetrodes fiber-coupled to a laser (**Figure 2C-H; Figure 2–supplement 1; Table 2**). Neurons that exhibited short-latency excitatory responses to optogenetic stimulation were positively identified as PT neurons (**Figure 2D**). All PT neurons exhibited action potential properties that are typical of cortical pyramidal neurons (Mitchell et al., 2007; Kaufman et al., 2010; Takahashi et al., 2015; Lohani et al., 2019) (**Figure 2–supplement 2**). The proportion of PT neurons that exhibited short-latency responses to optogenetic activation and the latency of those evoked responses (**Table 2**) were similar in Q175 and WT mice. Neurons that did not exhibit short-latency responses to optogenetic stimulation but were recorded on the same tetrode as identified PT neurons were also recorded. Unresponsive neurons that exhibited action potential properties that are typical of cortical pyramidal neurons (WT: 67%; Q175: 75%) were defined as putative layer V, IT neurons (**Figure 2D; Figure 2–supplement 2**). Consistent with previous studies, PT and putative IT neurons fired preferentially during the active component of cortical SWA (Steriade et al., 1993; Amzica and Steriade, 1998; Beltramo et al., 2013) (**Figure 2E**). During cortical SWA, the frequency and regularity (**Figure 2E-F; Table 2**) of PT and putative IT neuron activity were similar in Q175 and WT mice. In addition, pinch-evoked cortical desynchronization led to a similar reduction in PT and putative IT neuron activities in both Q175 and WT mice (**Figure 2G-H; Table 2**). Together, the EEG and single unit data suggest that 6-month-old Q175 and WT mice exhibit similar patterns and levels of motor cortical activity.

**Figure 2−supplement 1.**
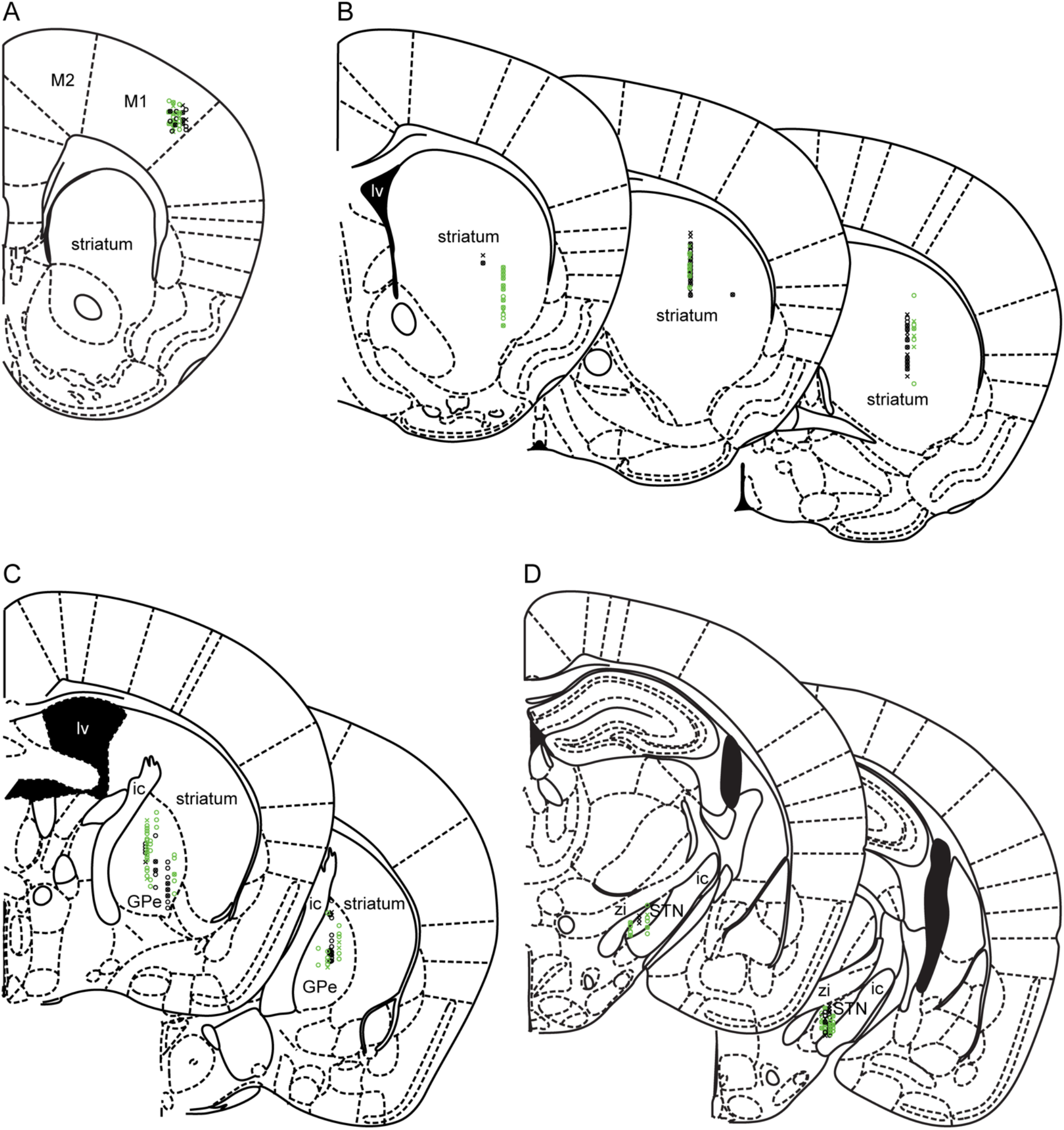
*In vivo* recording sites in the primary motor cortex, striatum, GPe, and STN. (**A-D**) The location of histologically confirmed electrode sites in which responsive (O) and non-responsive (X) neurons in WT (black) and Q175 (green) mice were recorded. (**A**) Recording sites in the primary motor cortex of PT-kj18-cre mice (M1, primary motor cortex; M2, secondary motor cortex). (**B**) Recording sites in the striatum of A2A-cre mice (lv, lateral ventricle). (**C**) Recording sites in the GPe of PV-cre mice (ic, internal capsule; rt, thalamic reticular nucleus). (**D**) Recording sites in the STN of PV-cre mice (zi, zona incerta).

**Figure 2−supplement 2.**
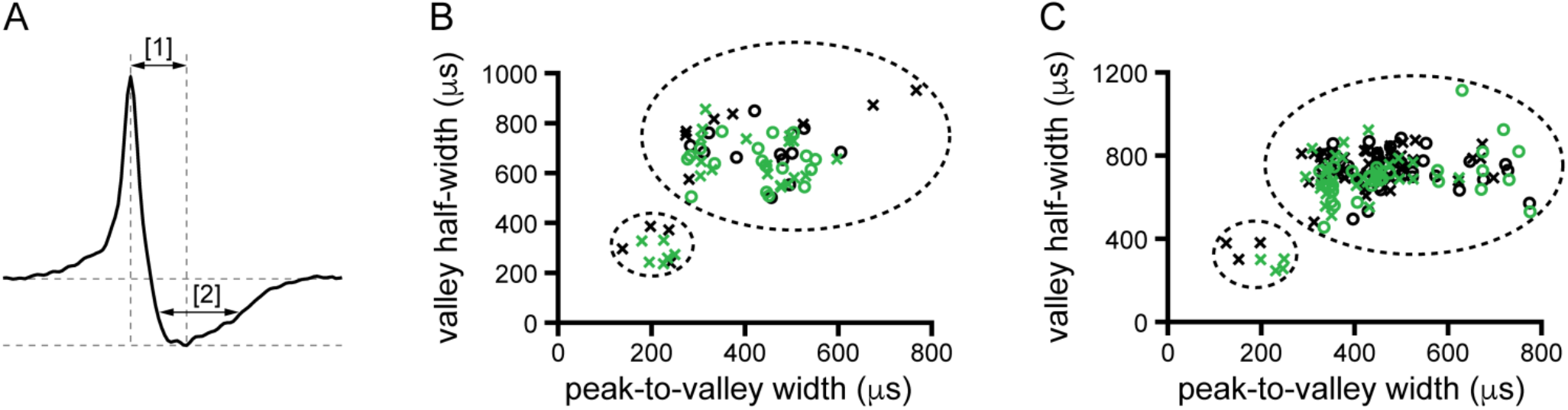
Classification of cortical and striatal neurons. (**A-C**) Action potential properties in optogenetically responsive (O) and non-responsive (X) neurons in WT (black) and Q175 (green) mice. Spike waveforms were measured from the average extracellular waveform of each unit and classified by their peak-to-valley width [1] and valley width at its half-maximum [2]. (**B, C**) Narrow and broad waveforms were used to discriminate putative interneurons and pyramidal cells in the cortex, respectively and putative interneurons and SPNs in the striatum, respectively (**C**).

**Table 2.**

| Figure/text | Measurement | Brain state | Genotype | n (mice/neurons) | Median [IQ range] | p-value (comparison; test) |
| --- | --- | --- | --- | --- | --- | --- |
| text | optogenetically responsive | - | WT | 5/14 | 63.6% | p = 0.42 (WT vs Q175; Fisher's Exact) |
|  |  |  | Q175 | 4/18 | 50.0% |  |
| text | latency of evoked response | - | WT | 5/14 | 5.0 [3.8-6.8] ms | p = 0.19 (WT vs Q175; MWU) |
|  |  |  | Q175 | 4/18 | 3.9 [3.3-5.4] ms |  |
| 2G | PT frequency | SWA | WT | 5/14 | 1.5 [0.60-2.1] Hz | p = 0.59 (SWA: WT vs Q175; MWU) |
|  |  |  | Q175 | 4/18 | 0.90 [0.20-1.8] Hz |  |
| 2G | IT frequency | SWA | WT | 4/8 | 2.5 [1.3-3.4] Hz | p = 0.42 (SWA: WT vs Q175; MWU) |
|  |  |  | Q175 | 4/18 | 1.1 [0.40-3.0] Hz |  |
| 2G | PT frequency | SWA | WT | 5/14 | 1.5 [0.60-2.1] Hz | p = 0.69 (SWA: PT vs IT; MWU) |
|  | IT frequency |  |  | 4/8 | 2.5 [1.3-3.4] Hz |  |
| 2G | PT frequency | SWA | Q175 | 4/18 | 0.90 [0.20-1.8] Hz | p = 0.45 (SWA: PT vs IT; MWU) |
|  | IT frequency |  |  | 4/18 | 1.1 [0.40-3.0] Hz |  |
| text | PT CV | SWA | WT | 5/12 | 1.3 [0.96-1.9] | p = 1.0 (SWA: WT vs Q175; MWU) |
|  |  |  | Q175 | 4/12 | 1.3 [0.85-2.0] |  |
| text | IT CV | SWA | WT | 4/8 | 1.4 [1.0-1.8] | p = 1.4 (SWA: WT vs Q175; MWU) |
|  |  |  | Q175 | 4/11 | 1.5 [1.3-2.3] |  |
| text | PT CV | SWA | WT | 5/12 | 1.3 [0.96-1.9] | p = 1.8 (SWA: PT vs IT; MWU) |
|  | IT CV |  |  | 4/8 | 1.4 [1.0-1.8] |  |
| text | PT CV | SWA | Q175 | 4/12 | 1.3 [0.85-2.0] | p = 1.2 (SWA: PT vs IT; MWU) |
|  | IT CV |  |  | 4/11 | 1.5 [1.3-2.3] |  |
| 2H | PT frequency | SWA | WT | 5/14 | 1.5 [0.60-2.1] Hz | p = 0.60 (SWA: WT vs Q175; MWU) |
|  |  |  | Q175 | 4/17 | 0.80 [0.20-2.0] Hz |  |
| 2H | PT frequency | ACT | WT | 5/14 | 0.0 [0.0-0.55] Hz | p = 0.020 (WT: SWA vs ACT; WSR) |
|  |  |  | Q175 | 4/17 | 0.0 [0.0-0.30] Hz | p = 4.0e-04 (Q175: SWA vs ACT; WSR) |
|  |  |  |  |  |  | p = 0.60 (ACT: WT vs Q175; MWU) |
| 2I | IT frequency | SWA | WT | 4/8 | 2.5 [1.3-3.4] Hz | p = 0.31 (SWA: WT vs Q175; MWU) |
|  |  |  | Q175 | 4/18 | 1.1 [0.40-3.0] Hz |  |
| 2I | IT frequency | ACT | WT | 4/8 | 0.0 [0.0-1.5] Hz | p = 0.062 (WT: SWA vs ACT; WSR) |
|  |  |  | Q175 | 4/18 | 0.20 [0.0-3.4] Hz | p = 0.45 (Q175: SWA vs ACT; WSR) |
|  |  |  |  |  |  | p = 0.54 (ACT: WT vs Q175; MWU) |
| text | PT frequency | ACT | WT | 5/14 | 0.0 [0.0-0.55] Hz | p = 0.86 (ACT: PT vs IT; MWU) |
|  | IT frequency |  |  | 4/8 | 0.0 [0.0-1.5] Hz |  |
| text | PT frequency | ACT | Q175 | 4/17 | 0.0 [0.0-0.30] Hz | p = 0.56 (ACT: PT vs IT; MWU) |
|  | IT frequency |  |  | 4/18 | 0.20 [0.0-3.4] Hz |  |

**Figure 2.**
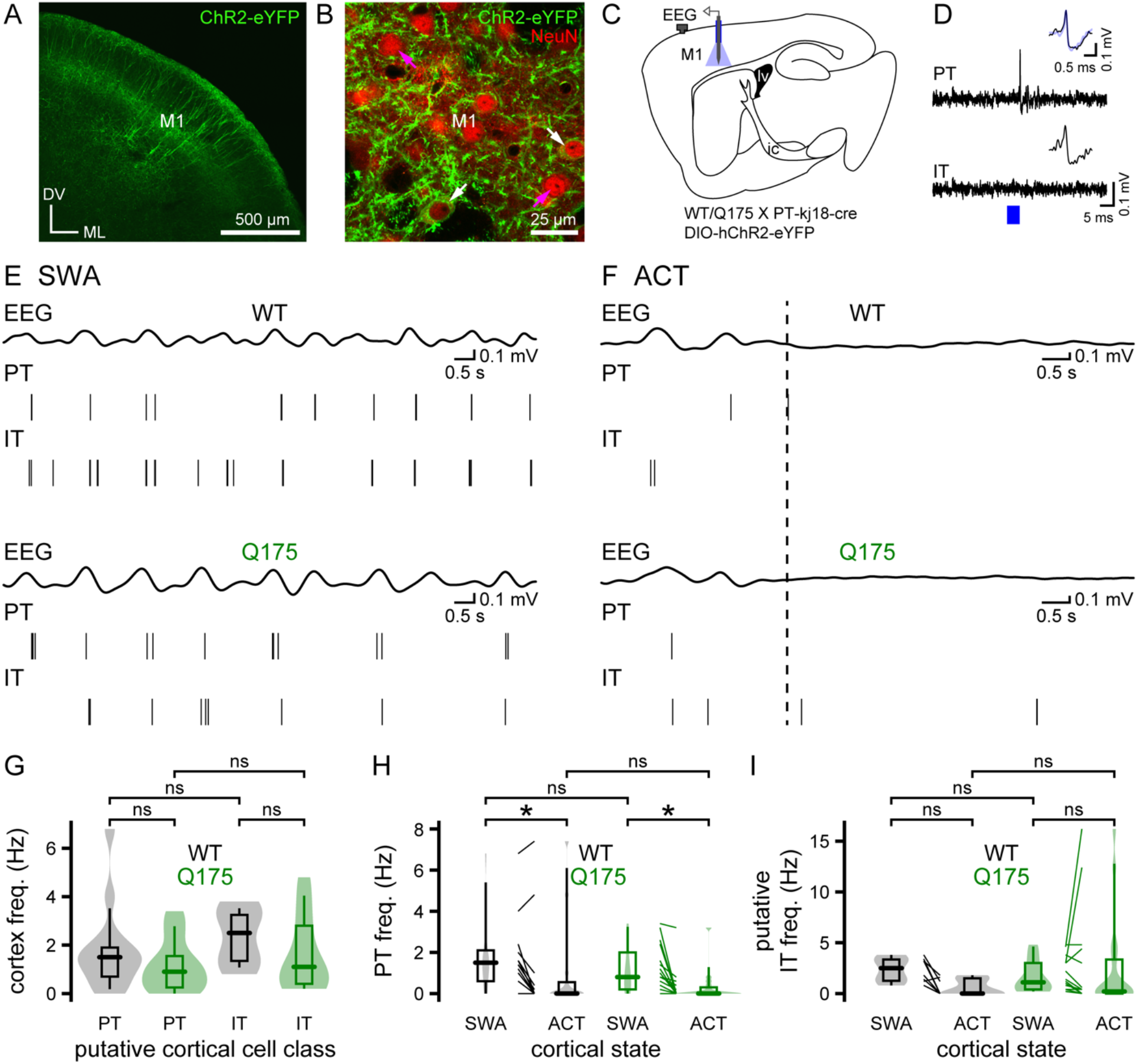
Motor cortical projection neuron activity is similar in Q175 and WT mice during both SWA and ACT. (**A-B**) Cre-dependent viral expression of ChR2(H134R)-eYFP (green) in layer 5 primary motor cortical PT neurons (M1, primary motor cortex; DV, dorsoventral axis; ML, mediolateral axis). (**B**) ChR2(H134R)-eYFP expression was present in a subset of layer V, presumably PT neurons (white arrows), and absent in putative layer V IT neurons and interneurons (magenta arrows). Cortical neurons were immunohistochemically labeled for the pan neuronal marker NeuN (red). (**C**) Schematic of experimental setup illustrating placement of the EEG screw electrode (gray) and optrode (blue). (**D**) An optogenetically stimulated PT neuron and an adjacent, unresponsive, putative IT neuron (blue, optogenetic stimulation; inset, spontaneous (black) and optogenetically-evoked (blue) action potentials **(E-I)**. The activities of identified PT and putative IT neurons in Q175 and age-matched WT mice were similar during cortical SWA and ACT. (**E, F**) Representative examples of concurrent EEG (band-pass filtered at 0.1-1.5 Hz) and cortical neuron activity during cortical SWA (**E**) and ACT (**F**). The frequency of firing of PT and putative IT neurons in both genotypes were similar during cortical SWA (**E**, examples; **G-I**, population data; PT: WT, n = 14 neurons; Q175, n = 18 neurons; IT: WT, n = 8 neurons; Q175, n = 18 neurons). (**F, H, I**) Hind paw pinch-evoked cortical ACT (dotted line) produced a similar reduction in PT and putative IT neuron activity in Q175 and WT mice (**F**, examples; **H-I**, population data). *, p < 0.05. ns, not significant.

### D2-SPNs are relatively hypoactive in Q175 mice

The striatum is largely composed of similar numbers of direct and indirect pathway striatal projection neurons (SPNs), which express D1 or D2 dopamine receptors, respectively (Albin et al., 1989; Gerfen et al., 1990). As their names suggest, direct pathway D1-SPNs directly innervate basal ganglia output neurons, whereas indirect pathway D2-SPNs regulate basal ganglia output indirectly via the GPe and STN (Mink and Thach, 1993; Maurice et al., 1999; Tachibana et al., 2008). Appropriate, cortical patterning of D1- and D2-SPN activity is key to the regulation of psychomotor function by the basal ganglia (Cui et al., 2013; Keeler et al., 2014; Nelson and Kreitzer, 2014; Tecuapetla et al., 2014; Sippy et al., 2015; Barbera et al., 2016; Lambot et al., 2016; Lemos et al., 2016; Tecuapetla et al., 2016; Klaus et al., 2019; LeBlanc et al., 2020). Although the striatum is a primary site of dysregulation and degeneration in HD and its models, precisely how the relative activities of D1-SPNs and D2-SPNs are perturbed *in vivo* is unknown. To address this question, ChR2(H134R)-eYFP was virally expressed in D2-SPNs through injection of an AAV carrying a cre-dependent expression construct into the striatum of Q175 and WT mice that had been crossed with a D2-SPN selective cre-driver line (A2A-cre). 2-3 weeks later the activities of striatal neurons in Q175 and WT mice were compared using silicon optrodes (**Figure 2–supplement 1**). Consistent with the successful targeting of D2-SPNs in both Q175 and WT mice, ChR2(H134R)-eYFP was expressed in a subset of striatal neurons projecting to the GPe but not to the substantia nigra *pars reticulata* (SNr) (**Figure 3A-C**). Therefore, optogenetic activation was used to positively identify D2-SPNs (**Figure 3D-E**). Consistent with their correct identification, all opto-tagged D2-SPNs exhibited action potential properties that are typical of SPNs rather than striatal interneurons (Berke et al., 2004; Gage et al., 2010; Cayzac et al., 2011; Kim et al., 2014; Shin et al., 2018) (**Figure 2–supplement 2**). The proportion of striatal neurons that exhibited short latency excitatory responses to optogenetic stimulation was similar in Q175 and WT mice (**Table 3**). The majority of adjacent neurons that were not optogenetically excited also exhibited action potential properties that are typical of striatal projection neurons (WT: 95%; Q175: 89%) (**Figure 2–supplement 2**) and were therefore classified as putative D1-SPNs. Consistent with direct cortical driving, identified and putative SPN activity was phase-locked to the active component of cortical SWA in both Q175 and WT mice (**Figure 3E**). In keeping with reduced axospinous synapse density, mEPSC amplitude, and dendritic excitability in D2-SPNs in HD mice, the frequency of D2-SPN firing was significantly lower in Q175 mice relative to WT controls (**Figure 3E-F; Table 3**). In contrast, the frequency of putative D1-SPN activity was similar in Q175 and WT mice during cortical SWA, despite reported reductions in glutamatergic synaptic excitation by some groups (Goodliffe et al., 2018) (**Figure 3F-G; Table 3**). Overall, during SWA the frequency of D1-SPN activity was greater than D2-SPN activity in Q175 but not WT mice (**Figure 3E-F; Table 3**). During pinch-evoked cortical ACT, the frequency of putative D1-SPN and D2-SPN activity in Q175 and WT mice declined (**Figure 3H-J; Table 3**). However, during cortical ACT the frequency of putative D1-SPN activity was greater in Q175 mice than WT mice (**Figure 3I-K; Table 3)**. Together, these data demonstrate that 1) during cortical SWA, D2-SPNs are hypoactive in Q175 mice relative to D2-SPNs in WT mice 2) during cortical SWA, D2-SPNs are hypoactive relative to putative D1-SPNs in Q175 but not WT mice.

**Table 3.**

| Figure/text | Measurement | Brain state | Genotype | n (mice/neurons) | Median [IQ range] | p-value (comparison; test) |
| --- | --- | --- | --- | --- | --- | --- |
| text | optogenetically responsive | - | WT | 3/30 | 36.1% | p = 0.13 (WT vs Q175; Fisher's Exact) |
|  |  |  | Q175 | 3/30 | 49.2% |  |
| text | latency of evoked response | - | WT | 3/30 | 6.8 [4.5-7.5] ms | p = 0.031 (WT vs Q175; MWU) |
|  |  |  | Q175 | 3/30 | 5.9 [4.4-6.5] ms |  |
| 3G | D2-SPN freq. | SWA | WT | 3/30 | 0.90 [0.35-2.4] Hz | p = 0.038 (SWA: WT vs Q175; MWU) |
|  |  |  | Q175 | 3/30 | 0.40 [0.0-0.80] Hz |  |
| 3G | D1-SPN freq. | SWA | WT | 3/53 | 0.80 [0.20-1.5] Hz | p = 1.0 (SWA: WT vs Q175; MWU) |
|  |  |  | Q175 | 3/31 | 0.60 [0.20-2.2] Hz |  |
| 3G | D2-SPN freq. | SWA | WT | 3/30 | 0.90 [0.35-2.4] Hz | p = 1.1 (SWA: D2 vs D1; MWU) |
|  | D1-SPN freq. |  |  | 3/53 | 0.80 [0.20-1.5] Hz |  |
| 3G | D2-SPN freq. | SWA | Q175 | 3/30 | 0.40 [0.0-0.80] Hz | p = 0.036 (SWA: D2 vs D1; MWU) |
|  | D1-SPN freq. |  |  | 3/31 | 0.60 [0.20-2.2] Hz |  |
| 3H | D2-SPN CV | SWA | WT | 3/20 | 1.6 [1.1-1.9] | p = 0.33 (SWA: WT vs Q175; MWU) |
|  |  |  | Q175 | 3/10 | 1.1 [0.68-1.7] |  |
| 3H | D1-SPN CV | SWA | WT | 3/35 | 1.3 [1.0-1.7] | p = 0.64 (SWA: WT vs Q175; MWU) |
|  |  |  | Q175 | 3/17 | 1.4 [1.1-2.1] |  |
| 3H | D2-SPN CV | SWA | WT | 3/20 | 1.6 [1.1-1.9] | p = 0.83 (SWA: D2 vs D1; MWU) |
|  | D1-SPN CV |  |  | 3/35 | 1.3 [1.0-1.7] |  |
| 3H | D2-SPN CV | SWA | Q175 | 3/10 | 1.1 [0.68-1.7] | p = 0.49 (SWA: D2 vs D1; MWU) |
|  | D1-SPN CV |  |  | 3/17 | 1.4 [1.1-2.1] |  |
| 3J | D2-SPN freq. | SWA | WT | 3/28 | 1.2 [0.40-2.4] Hz | p = 0.017 (SWA: WT vs Q175; MWU) |
|  |  |  | Q175 | 3/30 | 0.40 [0.0-0.80] Hz |  |
| 3J | D2-SPN freq. | ACT | WT | 3/28 | 0.0 [0.0-0.0] Hz | p = 1.1e-04 (WT: SWA vs ACT; WSR) |
|  |  |  | Q175 | 3/30 | 0.0 [0.0-0.05] Hz | p = 0.018 (Q175: SWA vs ACT; WSR) |
|  |  |  |  |  |  | p = 0.34 (ACT: WT vs Q175; MWU) |
| 3K | D1-SPN freq. | SWA | WT | 3/51 | 0.80 [0.20-1.6] Hz | p = 0.93 (SWA: WT vs Q175; MWU) |
|  |  |  | Q175 | 3/31 | 0.60 [0.20-2.2] Hz |  |
| 3K | D1-SPN freq. | ACT | WT | 3/51 | 0.0 [0.0-0.0] Hz | p = 3.6e-08 (WT: SWA vs ACT; WSR) |
|  |  |  | Q175 | 3/31 | 0.0 [0.0-1.2] Hz | p = 3.2e-03 (Q175: SWA vs ACT; WSR) |
|  |  |  |  |  |  | p = 6.9e-05 (ACT: WT vs Q175; MWU) |
| text | D2-SPN freq. | ACT | WT | 3/28 | 0.0 [0.0-0.0] Hz | p = 0.32 (ACT: D2 vs D1; MWU) |
|  | D1-SPN freq. |  |  | 3/51 | 0.0 [0.0-0.0] Hz |  |
| text | D2-SPN freq. | ACT | Q175 | 3/30 | 0.0 [0.0-0.05] Hz | p = 0.31 (ACT: D2 vs D1; MWU) |
|  | D1-SPN freq. |  |  | 3/31 | 0.0 [0.0-1.2] Hz |  |

**Figure 3.**
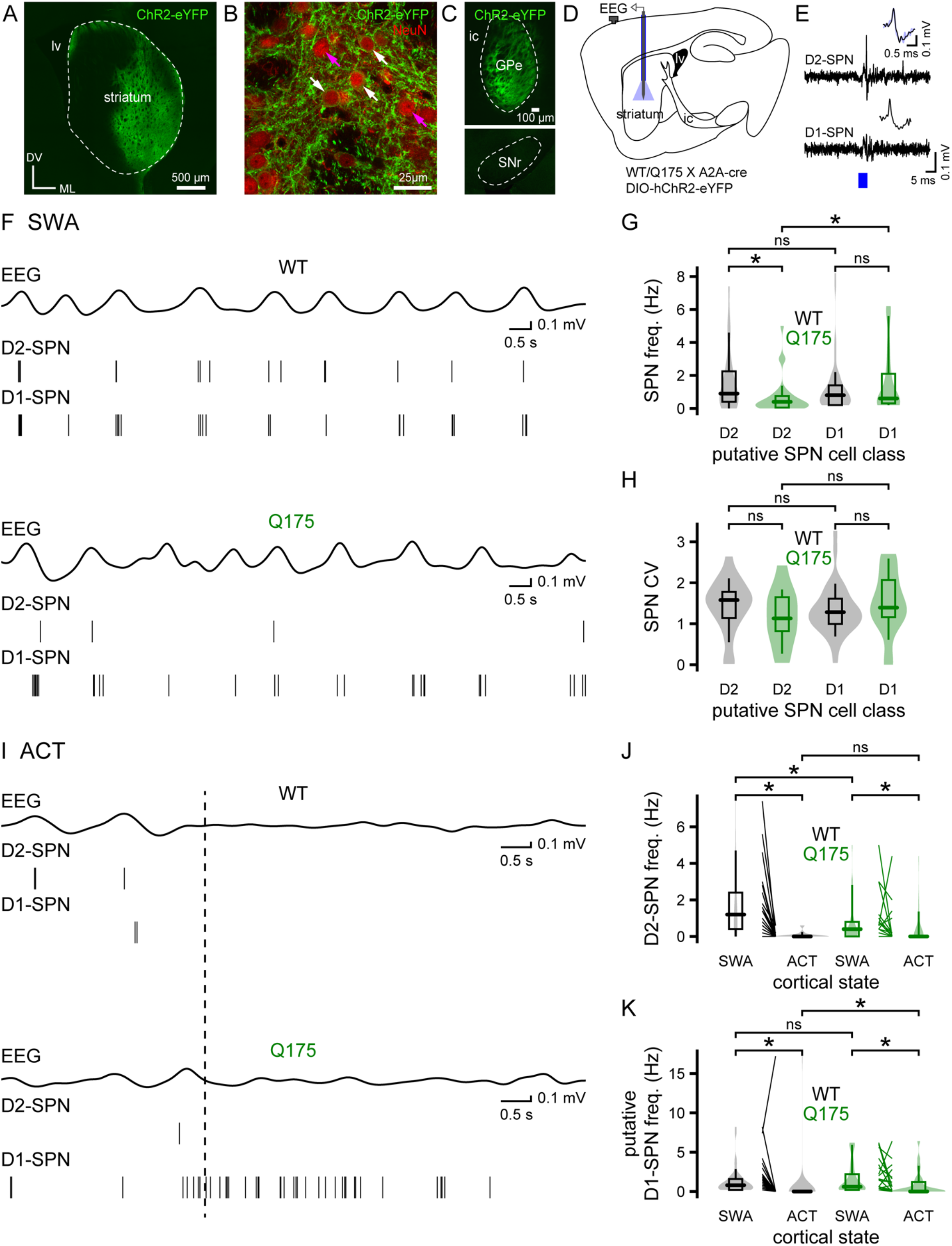
D2-SPNs are hypoactive in Q175 mice. (**A-C**) Viral-mediated, cre-dependent expression of hChR2(H134R)-eYFP (green) in D2-receptor expressing striatopallidal neurons in WT/Q175 X A2A-cre mice in the striatum (**A-B**; lv, lateral ventricle; DV, dorsoventral axis; ML, mediolateral axis) and GPe (**C**; ic, internal capsule; same orientation as **A**). (**B**) hChR2(H134R)-eYFP expression was present (white arrows)) or absent (magenta arrows) in striatal neurons that were co-immunoreactive for the pan neuronal marker NeuN (red). (**C**) Axon-terminal expression of hChR2(H134R)-eYFP in the GPe but not the SNr, consistent with selective expression in D2-but not D1-SPNs (ic, internal capsule). (**D**) Schematic of experimental set up, illustrating striatal placement of the optrode. (**E**) Example of optogenetic activation of a D2-but not a neighboring, putative D1-SPN (blue, optogenetic stimulation; inset, spontaneous (black) and optogenetically-evoked (blue) action potentials (**F-H**). The frequency of D2-SPN firing was lower in Q175 mice relative to age-matched WT mice during SWA (**F**, examples; **G-H**, population data). D2-SPNs also exhibited lower firing rates than putative D1-SPNs in Q175 but not WT mice (**G**, D2-SPNs: WT, n = 30 neurons; Q175, n = 30 neurons; putative D1-SPNs: WT, n = 53 neurons; Q175, n = 31 neurons). (**I-K**) Effect of pinch-evoked cortical ACT (dotted line) on both D2-SPNs and putative D1-SPNs in Q175 and WT mice (**I**, examples; **J, K**, population data; D2-SPNs: WT, n = 28 neurons; Q175, n = 30 neurons; putative D1-SPNs: WT, n = 51 neurons; Q175, n = 31 neurons). During cortical ACT the frequency of putative D1-SPN activity was higher in Q175 mice (**I, K**). *, p < 0.05. ns, not significant.

### Prototypic PV+ GPe neurons are hyperactive in Q175 mice

To determine whether there are changes in downstream GPe activity, we compared the *in vivo* firing of GPe neurons in Q175 and age-matched WT control mice. The GABAergic projection neurons of the GPe can be divided into two major types, so-called prototypic and arkypallidal neurons (Mallet et al., 2012; Mastro et al., 2014; Abdi et al., 2015; Dodson et al., 2015; Hernandez et al., 2015). Prototypic neurons comprise approximately three-quarters of all GPe neurons, are preferentially innervated by D2-SPNs, and innervate arkypallidal and downstream basal ganglia neurons (Mastro et al., 2014; Abdi et al., 2015; Dodson et al., 2015; Hernandez et al., 2015; Aristieta et al., 2020; Ketzef and Silberberg, 2020). A subset of prototypic neurons also innervates the striatum and/or cortex (Bevan et al., 1998; Mastro et al., 2014; Saunders et al., 2016; Abecassis et al., 2020). The majority of prototypic neurons express the calcium-binding protein parvalbumin (PV) (Mastro et al., 2014; Abdi et al., 2015; Dodson et al., 2015; Hernandez et al., 2015). In contrast, arkypallidal neurons comprise approximately one-fourth of GPe neurons, are preferentially innervated by D1-SPNs rather than D2-SPNs, innervate the striatum, rather than prototypic GPe neurons or the downstream basal ganglia, and express the transcription factor forkhead box protein 2 (FoxP2) but not PV (Mallet et al., 2012; Mastro et al., 2014; Abdi et al., 2015; Dodson et al., 2015; Hernandez et al., 2015; Aristieta et al., 2020; Ketzef and Silberberg, 2020). Although D2-SPNs do not directly target arkypallidal neurons, D2-SPNs can indirectly and powerfully regulate arkypallidal activity via their effects on prototypic GPe neurons (Aristieta et al., 2020; Ketzef and Silberberg, 2020).

Because the majority of GPe neurons fire tonically, we used optogenetic silencing to identify and manipulate these cells *in vivo*. The inhibitory opsin Arch-GFP was virally expressed in prototypic PV+ GPe neurons through injection of an AAV vector carrying a cre-dependent expression construct into the GPe of Q175 and WT mice that had been crossed with a PV-cre driver line (**Figure 4A-D**). Consistent with the selective expression of Arch-GFP in prototypic GPe neurons, the majority of Arch-GFP-expressing GPe neurons were immunoreactive for PV and all were immunonegative for FoxP2 (**Figure 4B**). In addition, Arch-GFP was expressed in the axon terminal fields of PV+ GPe neurons in the downstream basal ganglia, including the STN, consistent with their prototypic identity (**Figure 4C**). Optogenetic activation of Arch-GFP inhibited a similar proportion of neurons in the GPe of Q175 and WT mice (**Figure 2–supplement 1; Table 4**). As previously reported (Abdi et al., 2015; Kovaleski et al., 2020), PV+ GPe neurons discharged tonically during cortical SWA (**Figure 4E**). However, the frequency of prototypic PV+ GPe neuron activity was significantly greater in Q175 mice relative to WT controls (**Figure 4E-F; Table 4**). In Q175 mice the median PV+ GPe neuron activity was approximately double that in WT mice (**Figure 4E-F; Table 4**). While there was no significant difference in the overall regularity of PV+ GPe neuron firing in Q175 and WT mice, as assessed from the CV of the interspike interval (**Figure 4G; Table 4**), PV+ GPe neuron firing was elevated during the inactive relative to the active component of the EEG in Q175 mice (**Figure 4H-I; Table 4**). During pinch-evoked cortical ACT, the firing rate of PV+ GPe neurons increased in both genotypes (**Figure 4J-K; Table 4**). However, PV+ GPe neurons remained hyperactive in Q175 mice relative to WT controls (**Figure 4J-K; Table S4**). Together, these data demonstrate that prototypic PV+ GPe neuron activity is greatly elevated during both cortical SWA and ACT and is relatively antiphasic to cortical SWA in Q175 mice.

**Table 4.**
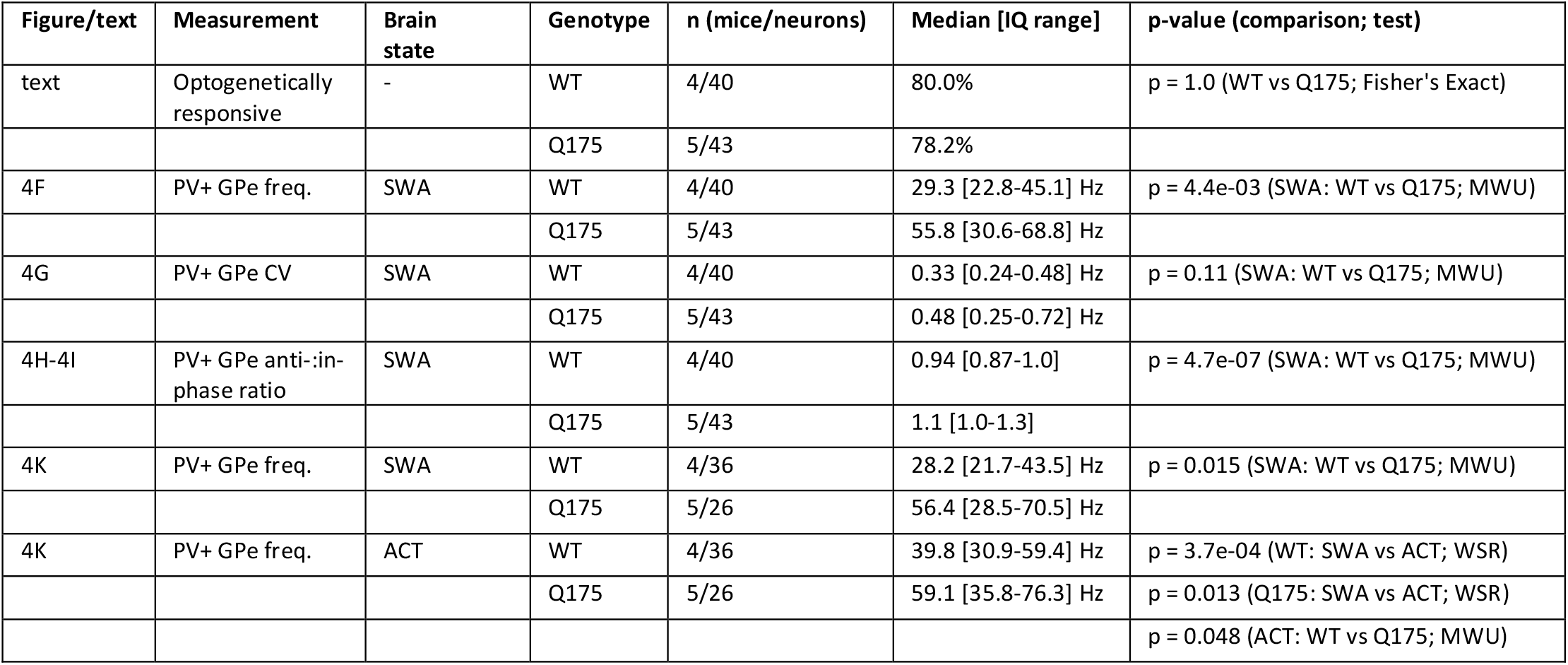

**Figure 4.**
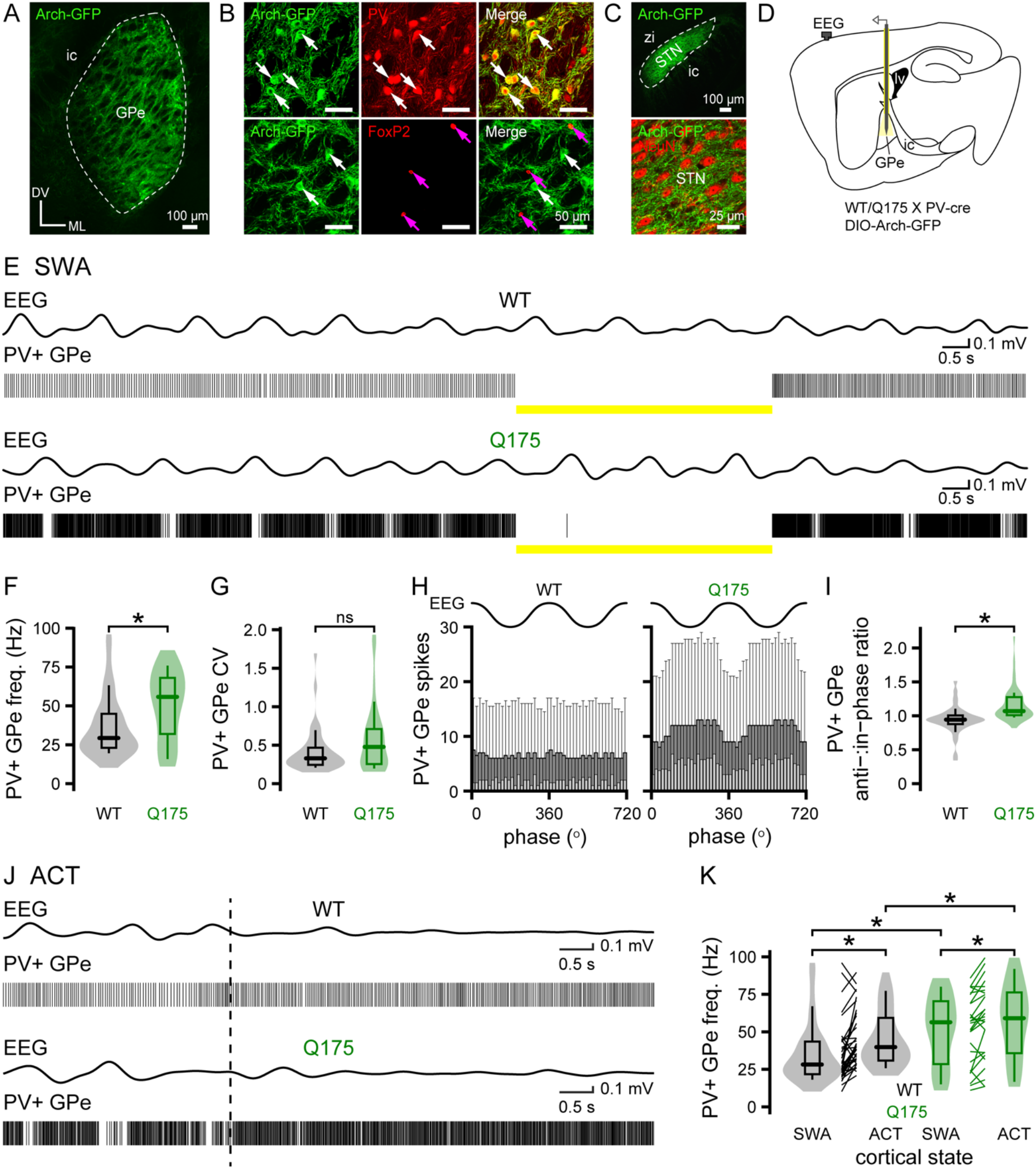
In Q175 mice PV+ GPe neurons are hyperactive and their firing is relatively antiphasic to cortical SWA. (**A-C**) Viral cre-dependent expression of Arch-GFP (green) in prototypic PV+ GPe neurons in Q175/WT X PV-cre mice. (**A**) Representative confocal micrograph of the GPe (ic, internal capsule; DV, dorsoventral axis; ML, mediolateral axis). (**B**) Arch-GFP expression was present in PV-immunoreactive GPe neurons (white arrows; upper panel) and FoxP2-immunonegative neurons (white arrows; lower panel) and absent in FoxP2-immunoreactive arkypallidal GPe neurons (magenta arrows; lower panel). (**C**) Arch-GFP-labeled axon terminal fields in the STN (upper panel; zi, zona incerta; same orientation as **A**) and in the vicinity of NeuN-immunoreactive STN neurons (red; lower panel). (**D**) Schematic of the experimental set up illustrating optrode placement in the GPe. (**E-K**) The firing of PV+ GPe neurons was altered in Q175 mice. (**E**) Representative examples of concurrent EEG (band-pass filtered at 0.1-1.5 Hz) and PV+ GPe neuron activity in the absence and presence of optogenetic inhibition (yellow bar). (**E-G**) During cortical SWA the median frequency of PV+ GPe firing in Q175 mice was approximately double that in WT mice but the variability of firing was unaltered (**E**, representative examples; **F, G**, population data; WT, n = 40 neurons; Q175, n = 43 neurons). (**H, I**) In Q175 mice the firing of PV+ GPe neurons was also relatively antiphasic to cortical SWA (**H**, population spike phase histograms; **I**, population box and violin plots). (**J, K**) Pinch-evoked cortical ACT (hind paw pinch, dotted line) increased PV+ GPe neuron firing in both genotypes. However, the frequency of PV+ GPe neuron discharge remained elevated in Q175 mice relative to WT mice (**J**, representative examples; **K**, population data; WT, n = 36 neurons; Q175, n = 26 neurons). *, p < 0.05. ns, not significant.

### The abnormal hypoactivity of PV-GPe neurons in Q175 mice is reversed by optogenetic inhibition of PV+ GPe neurons

We next analyzed the activity of PV-GPe, putative arkypallidal neurons, i.e. GPe neurons that were not directly inhibited during optogenetic activation of Arch-GFP in PV-cre mice (**Figure 2–supplement 1**). During cortical SWA and ACT, PV-GPe neurons were less active than PV+ GPe neurons in both WT and Q175 mice, consistent with previous reports in WT rodents (**Figure 5A-B; Table 5**) (Abdi et al., 2015; Dodson et al., 2015; Mallet et al., 2016). PV-GPe neurons also discharged more slowly in Q175 than WT mice (**Figure 5A-B; Table 5**). Recent studies suggest that prototypic GPe neurons inhibit arkypallidal neurons through powerful, local connections, whereas arkypallidal neuron innervation of prototypic GP neurons is negligible (Aristieta et al., 2020; Ketzef and Silberberg, 2020). If elevated inhibition emanating from hyperactive prototypic PV+ GPe neurons is responsible for the hypoactivity of PV-GPe, putative arkypallidal neurons in Q175 mice, then PV-GPe neuron activity should be rescued by optogenetic inhibition of PV+ GPe neurons. Optogenetic silencing of PV+ GPe neurons during cortical SWA disinhibited PV-GPe neuron activity in both WT and Q175 mice, confirming that PV+ GPe neurons actively inhibit PV-GPe neuron activity *in vivo* (**Figure 5A, D; Table 5**). Furthermore, during optogenetic inhibition of PV+ GPe neurons, the activities of PV-GPe neurons in WT and Q175 mice were no longer significantly different (**Figure 5D-E; Table 5**), arguing that increased inhibition from hyperactive PV+ GPe neurons was indeed responsible for the relative hypoactivity of putative arkypallidal neurons in Q175 mice. No consistent phase differences in the activity of PV-GPe neurons with or without optogenetic inhibition of PV+ GPe neurons were observed in WT and Q175 mice (**Figure 5E-F; Table 5**).

**Table 5.**
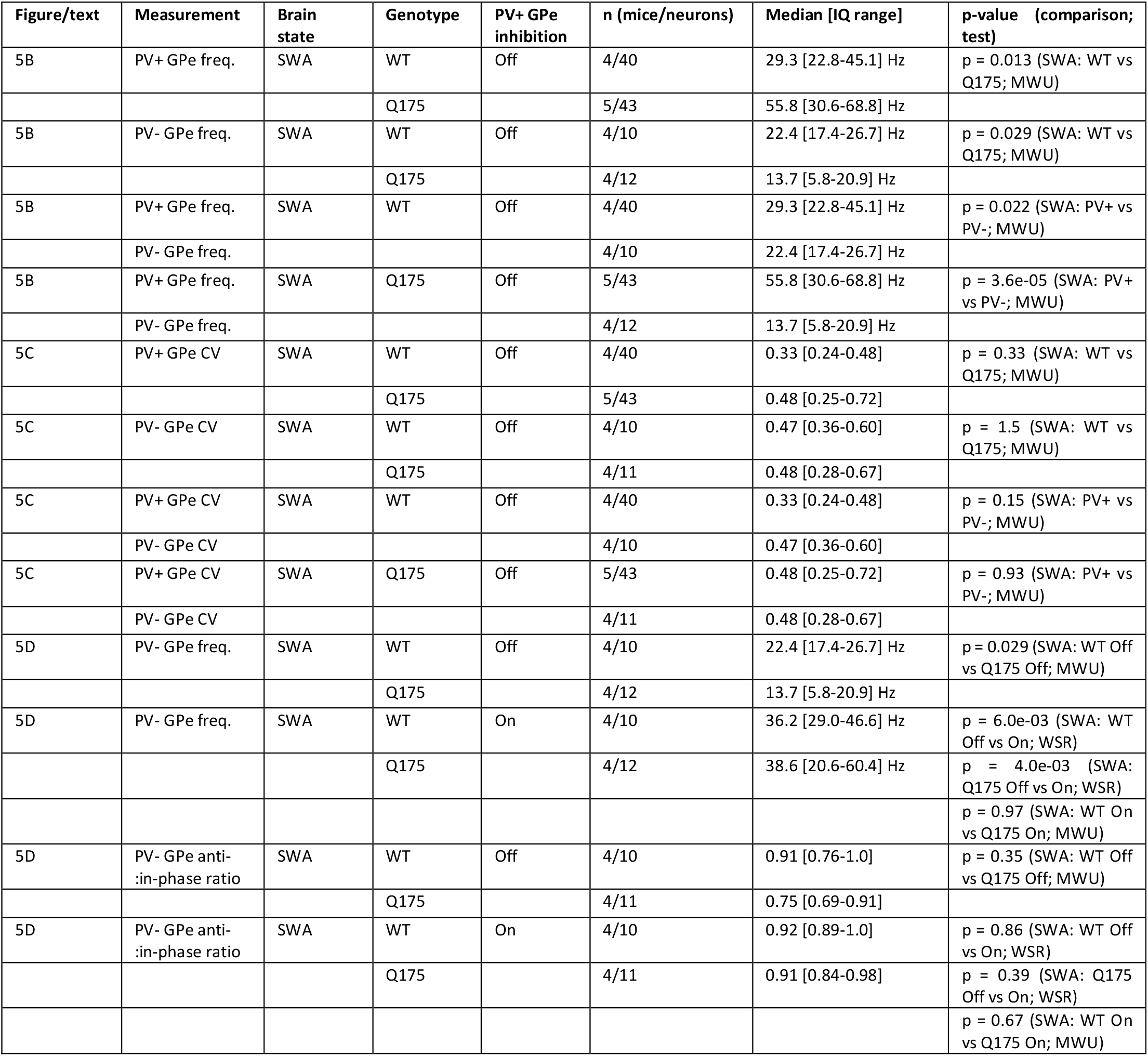

| Figure/text | Measurement | Brain state | Genotype | PV+ GPe inhibition | n (mice/neurons) | Median [IQ range] | p-value (comparison; test) |
| --- | --- | --- | --- | --- | --- | --- | --- |
| 5B | PV+ GPe freq. | SWA | WT | Off | 4/40 | 29.3 [22.8-45.1] Hz | p = 0.013 (SWA: WT vs Q175; MWU) |
|  |  |  | Q175 |  | 5/43 | 55.8 [30.6-68.8] Hz |  |
| 5B | PV- GPe freq. | SWA | WT | Off | 4/10 | 22.4 [17.4-26.7] Hz | p = 0.029 (SWA: WT vs Q175; MWU) |
|  |  |  | Q175 |  | 4/12 | 13.7 [5.8-20.9] Hz |  |
| 5B | PV+ GPe freq. | SWA | WT | Off | 4/40 | 29.3 [22.8-45.1] Hz | p = 0.022 (SWA: PV+ vs PV-; MWU) |
|  | PV- GPe freq. |  |  |  | 4/10 | 22.4 [17.4-26.7] Hz |  |
| 5B | PV+ GPe freq. | SWA | Q175 | Off | 5/43 | 55.8 [30.6-68.8] Hz | p = 3.6e-05 (SWA: PV+ vs PV-; MWU) |
|  | PV- GPe freq. |  |  |  | 4/12 | 13.7 [5.8-20.9] Hz |  |
| 5C | PV+ GPe CV | SWA | WT | Off | 4/40 | 0.33 [0.24-0.48] | p = 0.33 (SWA: WT vs Q175; MWU) |
|  |  |  | Q175 |  | 5/43 | 0.48 [0.25-0.72] |  |
| 5C | PV- GPe CV | SWA | WT | Off | 4/10 | 0.47 [0.36-0.60] | p = 1.5 (SWA: WT vs Q175; MWU) |
|  |  |  | Q175 |  | 4/11 | 0.48 [0.28-0.67] |  |
| 5C | PV+ GPe CV | SWA | WT | Off | 4/40 | 0.33 [0.24-0.48] | p = 0.15 (SWA: PV+ vs PV-; MWU) |
|  | PV- GPe CV |  |  |  | 4/10 | 0.47 [0.36-0.60] |  |
| 5C | PV+ GPe CV | SWA | Q175 | Off | 5/43 | 0.48 [0.25-0.72] | p = 0.93 (SWA: PV+ vs PV-; MWU) |
|  | PV- GPe CV |  |  |  | 4/11 | 0.48 [0.28-0.67] |  |
| 5D | PV- GPe freq. | SWA | WT | Off | 4/10 | 22.4 [17.4-26.7] Hz | p = 0.029 (SWA: WT Off vs Q175 Off; MWU) |
|  |  |  | Q175 |  | 4/12 | 13.7 [5.8-20.9] Hz |  |
| 5D | PV- GPe freq. | SWA | WT | On | 4/10 | 36.2 [29.0-46.6] Hz | p = 6.0e-03 (SWA: WT Off vs On; WSR) |
|  |  |  | Q175 |  | 4/12 | 38.6 [20.6-60.4] Hz | p = 4.0e-03 (SWA: Q175 Off vs On; WSR) |
|  |  |  |  |  |  |  | p = 0.97 (SWA: WT On vs Q175 On; MWU) |
| 5D | PV- GPe anti-in-phase ratio | SWA | WT | Off | 4/10 | 0.91 [0.76-1.0] | p = 0.35 (SWA: WT Off vs Q175 Off; MWU) |
|  |  |  | Q175 |  | 4/11 | 0.75 [0.69-0.91] |  |
| 5D | PV- GPe anti-in-phase ratio | SWA | WT | On | 4/10 | 0.92 [0.89-1.0] | p = 0.86 (SWA: WT Off vs On; WSR) |
|  |  |  | Q175 |  | 4/11 | 0.91 [0.84-0.98] | p = 0.39 (SWA: Q175 Off vs On; WSR) |
|  |  |  |  |  |  |  | p = 0.67 (SWA: WT On vs Q175 On; MWU) |

**Figure 5.**
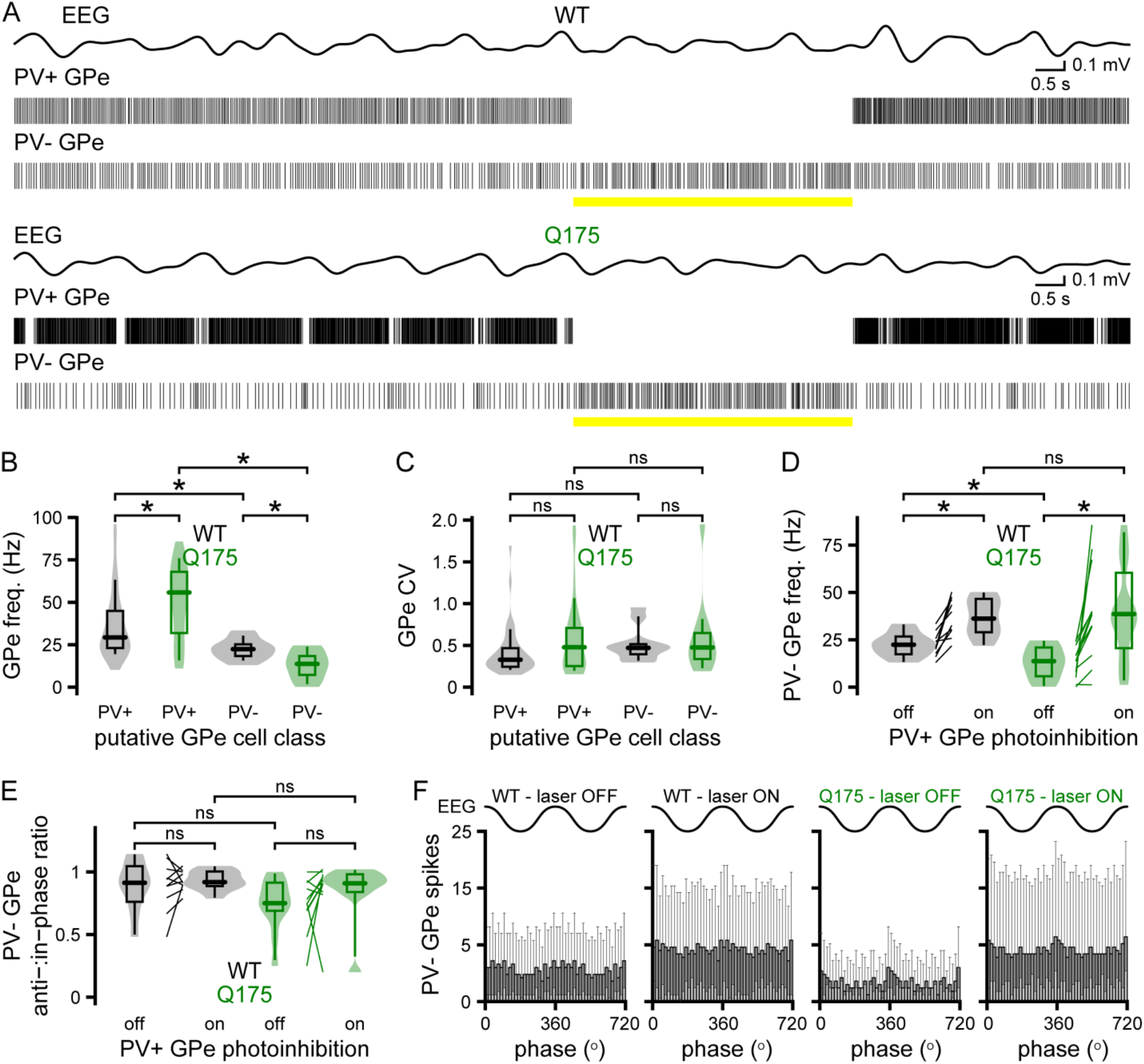
The hypoactivity of PV-GPe neurons in Q175 mice is alleviated by optogenetic inhibition of hyperactive PV+ GPe neurons. (**A**) Representative examples of concurrent EEG (band-pass filtered at 0.1-1.5 Hz) and GPe neuronal activity in Q175/WT X PV-cre mice during optogenetic inhibition (yellow bar) of PV+ GPe neurons. (**A-C**) During cortical SWA, PV-GPe neurons were less active than PV+ GPe neurons in both WT and Q175 mice. The frequency of PV-GPe neuron activity was also lower in Q175 mice than in WT mice (**A**, representative examples; **B, C**, population data; WT, n = 10 neurons; Q175, n = 12 neurons). Optogenetic inhibition of PV+ GPe neurons disinhibited PV-GPe neurons in WT and Q175 mice and eliminated the difference in firing frequencies in the two genotypes, arguing that GABAergic inhibition emanating from abnormally hyperactive PV+ GPe neurons is responsible for the relative hypoactivity of GPe PV-neurons in Q175 mice (**D**, population data). (**E, F**) Relative to cortical SWA, there were no phase differences in PV-GPe neuron activity in WT and Q175 mice, with or without optogenetic inhibition of PV+ GPe neurons. *, p < 0.05. ns, not significant.

### The autonomous activity of PV+ GPe neurons but not PV-GPe neurons is increased in brain slices from Q175 mice

The high rates of discharge of extrastriatal basal ganglia neurons *in vivo* are generated in part by their intrinsic autonomous activity (Wilson, 2013). Alterations in the autonomous firing of PV+ and PV-GPe neurons could therefore contribute to their abnormal *in vivo* activity in Q175 mice. To determine whether the autonomous firing of GPe neurons is altered in Q175 mice, an AAV vector carrying an eGFP-dependent expression construct was injected into the GPe of WT or Q175 PV-cre mice. 2-3 weeks later brain slices were prepared and visually guided somatic patch clamp recordings of GPe neurons were conducted. Expression of eGFP was used to identify prototypic PV+ GPe neurons. PV-GPe, putative arkypallidal neurons in the vicinity of PV+ GPe neurons were identified by their absence of eGFP expression. GPe neurons were recorded in the loose-seal, cell-attached, current-clamp configuration and recordings were made in the presence of AMPA, NMDA, GABA_A_, and GABA_B_ receptor antagonists to minimize the impact of synaptic inputs on GPe neuron activity. As for previous studies (Abdi et al., 2015; Hernandez et al., 2015), we found that the frequency and regularity of autonomous firing in prototypic PV+ GPe neurons were greater than in PV-GPe, putative arkypallidal neurons in WT mice (**Figure 6A-D; Table 6**). Furthermore, in Q175 mice the frequency and regularity of autonomous PV+ GPe neuron activity were significantly greater than in WT mice (**Figure 6A, C, D; Table 6**). In contrast, there was no difference in the frequency or regularity of autonomous PV-GPe neuron firing in Q175 and WT mice (**Figure 6B-D; Table 6**). Together, these *ex vivo* data demonstrate that the autonomous firing of prototypic PV+ GPe neurons is upregulated in Q175 mice and that this alteration may contribute to the hyperactivity of these cells *in vivo*. In contrast, the similarity of autonomous PV-GPe neuron firing in WT and Q175 mice further supports the conclusion that excessive inhibition of PV-GPe neurons emanating from hyperactive PV+ GPe neurons is responsible for the hypoactivity of PV-GPe, putative arkypallidal neurons in Q175 mice *in vivo*.

**Table 6.**

| Figure | Measurement | Genotype | n (mice/neurons) | Median [IQ range] | p-value (comparison; test) |
| --- | --- | --- | --- | --- | --- |
| 6C | PV+ GPe freq. | WT | 7/82 | 20.4 [14.8-28.8] Hz | p = 3.6e-03 (WT vs Q175; MWU) |
|  |  | Q175 | 8/73 | 25.9 [18.5-33.8] Hz |  |
| 6C | PV- GPe freq. | WT | 3/27 | 9.9 [6.6-12.8] Hz | p = 0.99 (WT vs Q175; MWU) |
|  |  | Q175 | 3/16 | 9.9 [5.3-13.0] Hz |  |
| 6C | PV+ GPe freq. | WT | 7/82 | 20.4 [14.8-28.8] Hz | p = 1.9e-10 (PV+ vs PV-; MWU) |
|  | PV- GPe freq. |  | 3/27 | 9.9 [6.6-12.8] Hz |  |
| 6C | PV+ GPe freq. | Q175 | 8/73 | 25.9 [18.5-33.8] Hz | p = 3.0e-08 (PV+ vs PV-; MWU) |
|  | PV- GPe freq. |  | 3/16 | 9.9 [5.3-13.0] Hz |  |
| 6D | PV+ GPe CV | WT | 7/82 | 0.13 [0.10-0.19] | p = 1.2e-03 (WT vs Q175; MWU) |
|  |  | Q175 | 8/73 | 0.10 [0.082-0.14] |  |
| 6D | PV- GPe CV | WT | 3/27 | 0.19 [0.14-0.37] | p = 0.79 (WT vs Q175; MWU) |
|  |  | Q175 | 3/16 | 0.22 [0.17-0.29] |  |
| 6D | PV+ GPe CV | WT | 7/82 | 0.13 [0.10-0.19] | p = 4.4e-04 (PV+ vs PV-; MWU) |
|  | PV- GPe CV |  | 3/27 | 0.19 [0.14-0.37] |  |
| 6D | PV+ GPe CV | Q175 | 8/73 | 0.10 [0.082-0.14] | p = 2.1e-06 (PV+ vs PV-; MWU) |
|  | PV- GPe CV |  | 3/16 | 0.22 [0.17-0.29] |  |

**Figure 6.**
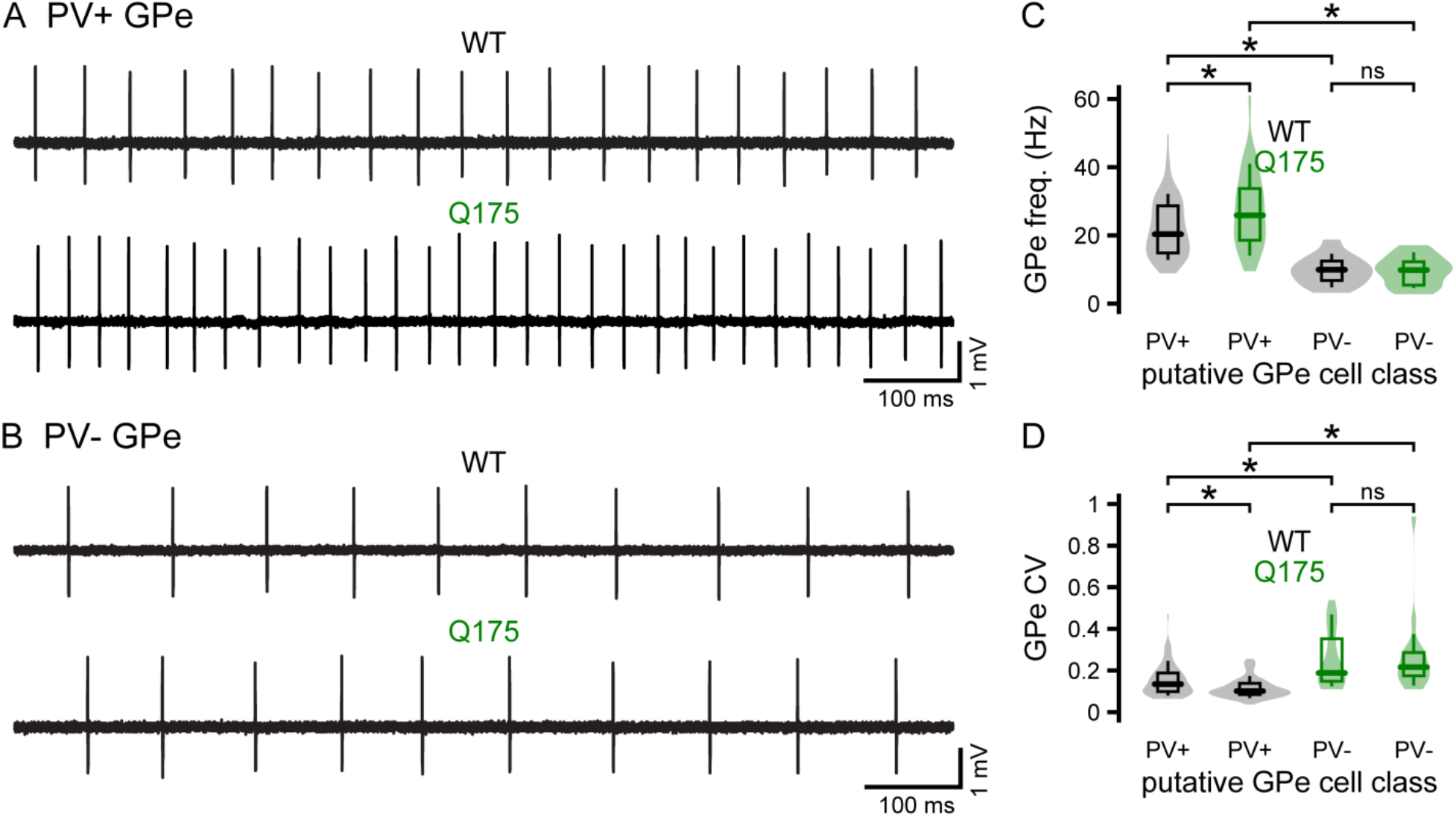
The autonomous activity of PV+ GPe neurons is elevated in Q175 mice. (**A, B**) Representative examples of autonomous GPe neuron activity recorded in the loose-seal, cell-attached, current-clamp configuration in brain slices from Q175/WT X PV-cre mice. (**A, C, D**) The frequency and regularity of autonomous PV+ GPe neuron activity were greater in slices from Q175 mice (**A**, representative examples; **C, D**, population data; WT, n = 82 neurons; Q175, n = 73 neurons). (**B-D**) The frequency and regularity of autonomous PV-GPe neuron activity were similar in Q175 and WT mice (**B**, representative examples; **C, D**, population data; WT, n = 27 neurons; Q175, n = 16 neurons). *, p < 0.05. ns, not significant.

### The hypoactivity of the STN in Q175 mice is partially alleviated by optogenetic inhibition of prototypic PV+ GPe neurons

The glutamatergic STN is a key component of the indirect pathway and forms a reciprocally connected network with the GPe (Mink and Thach, 1993; Maurice et al., 1999; Plenz and Kital, 1999; Nambu et al., 2002; Tachibana et al., 2008). Previous research has demonstrated that autonomous STN activity is downregulated in HD mice (Atherton et al., 2016). Furthermore, we demonstrate here that GABAergic prototypic PV+ GPe neurons, which potently inhibit STN activity through their activation of postsynaptic GABA_A_ and GABA_B_ receptors (Bevan et al., 2002; Hallworth and Bevan, 2005; Baufreton et al., 2009; Atherton et al., 2013; Kovaleski et al., 2020), are hyperactive in Q175 mice *in vivo*. Given the loss of intrinsic STN activity and increased frequency of GABAergic GPe-STN transmission, we predicted that STN neurons will be hypoactive in Q175 mice *in vivo* compared to WT. To test this prediction, we recorded the activity of STN neurons with silicon tetrodes during cortical SWA and ACT in WT and Q175 mice crossed with PV-cre mice (**Figure 2–supplement 1**). To isolate the effect of PV+ GPe neuron inhibition on the STN, we virally expressed Arch-GFP in PV+ GPe neurons and inhibited their activity optogenetically, as described above. Similar to previous studies, we found that STN neurons fired in phase with cortical SWA (Magill et al., 2000, 2001; Mallet et al., 2008a; Callahan and Abercrombie, 2015b, a; Kovaleski et al., 2020). However, the frequency (but not the regularity) of STN activity was reduced in Q175 mice (**Figure 7A-D; Table 7**). The proportion of spikes generated during the inactive component of cortical SWA relative to those during the active component was similar in Q175 and WT mice (**Figure 7E-F; Table 7**). Pinch-evoked cortical ACT increased the frequency of STN activity in both WT and Q175 mice (**Figure 7G-H; Table 7**). However, the frequency of STN activity remained lower in Q175 mice relative to that in WT mice during cortical ACT (**Figure 7G-H; Table 7**).

**Table 7.**

| Figure/text | Measurement | Brain state | Genotype | n (mice/neurons) | Median [IQ range] | p-value (comparison; test) |
| --- | --- | --- | --- | --- | --- | --- |
| 7C | STN freq. | SWA | WT | 4/30 | 7.6 [4.6-13.3] Hz | p = 7.0e-04 (SWA: WT vs Q175; MWU) |
|  |  |  | Q175 | 5/61 | 4.4 [2.5-7.6] Hz |  |
| 7D | STN CV | SWA | WT | 4/30 | 1.8 [1.3-2.3] | p = 0.41 (SWA: WT vs Q175; MWU) |
|  |  |  | Q175 | 5/60 | 1.7 [1.2-2.4] |  |
| 7E-7F | STN anti-in-phase ratio | SWA | WT | 4/30 | 0.32 [0.11-0.53] | p = 0.25 (SWA: WT vs Q175; MWU) |
|  |  |  | Q175 | 5/61 | 0.25 [0.064-0.38] |  |
| 7H | STN freq. | SWA | WT | 4/26 | 7.2 [4.4-13.0] Hz | p = 5.8e-03 (SWA: WT vs Q175; MWU) |
|  |  |  | Q175 | 5/51 | 4.6 [2.4-7.2] Hz |  |
| 7H | STN freq. | ACT | WT | 4/26 | 17.0 [9.3-21.7] Hz | p = 4.0e-04 (WT: SWA vs ACT; WSR) |
|  |  |  | Q175 | 5/51 | 7.8 [4.0-13.6] Hz | p = 3.0e-04 (Q175: SWA vs ACT; WSR) |
|  |  |  |  |  |  | p = 3.2e-04 (ACT: WT vs Q175; MWU) |

**Figure 7.**
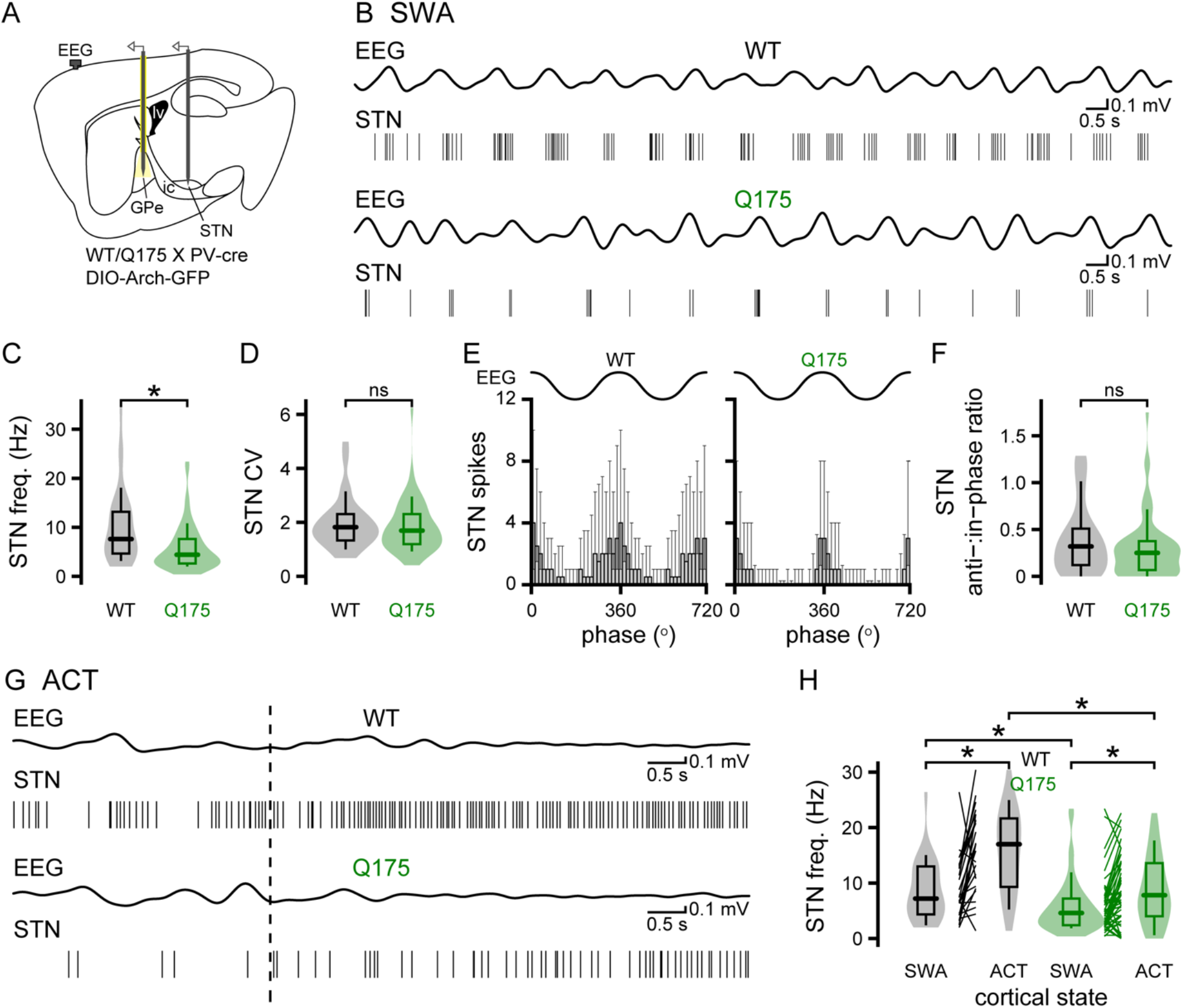
STN neurons are hypoactive in Q175 mice. (**A-D**) During cortical SWA, the frequency (but not the regularity) of STN activity was lower in Q175 mice (**A**, schematic of experimental set up, illustrating placement of an optrode in the GPe and a tetrode array in the STN; **B**, representative examples of concurrent 0.1-1.5 Hz band-pass filtered EEG and STN unit activity; **C, D**, population data; WT, n = 30 neurons; Q175, n = 61 neurons). (**E, F**) STN activity was similarly phase-locked to the active relative to the inactive component of cortical SWA in Q175 and WT mice (**E**, population spike phase histograms; **F**, population data). (**G, H**) Pinch-evoked cortical ACT (hind paw pinch, dotted line) increased STN neuron activity in both genotypes. However, during cortical ACT, STN neurons remained hypoactive in Q175 mice relative to WT (**G**, representative examples; **H**, population data; WT, n = 26 neurons; Q175, n = 51 neurons). *, p < 0.05. ns, not significant.

To determine how PV+ GPe neurons regulate STN activity, the response of STN neurons to optogenetic inhibition of PV+ GPe neurons was recorded. The proportion of STN neurons that responded to inhibition of PV+ GPe neurons was similar in WT and Q175 mice (**Table 8**). Optogenetic inhibition of PV+ GPe neurons increased the frequency and regularity of STN activity in both WT and Q175 mice (**Figure 8A-C; Table 8**). However, during optogenetic inhibition, STN neurons remained less active in Q175 mice during cortical SWA (**Figure 8A-C; Table 8**) and ACT (**Table 8**). The ratio of anti- to in-phase STN activity was also significantly elevated by inhibition of PV+ GPe neurons in Q175 mice (**Figure 8D-E; Table 8**). Together, with data from this and previous studies, these findings argue that both excessive inhibition arising from hyperactive PV+ GPe neurons *and* autonomous firing deficits contribute to the relative hypoactivity of STN neurons in Q175 mice.

**Table 8.**
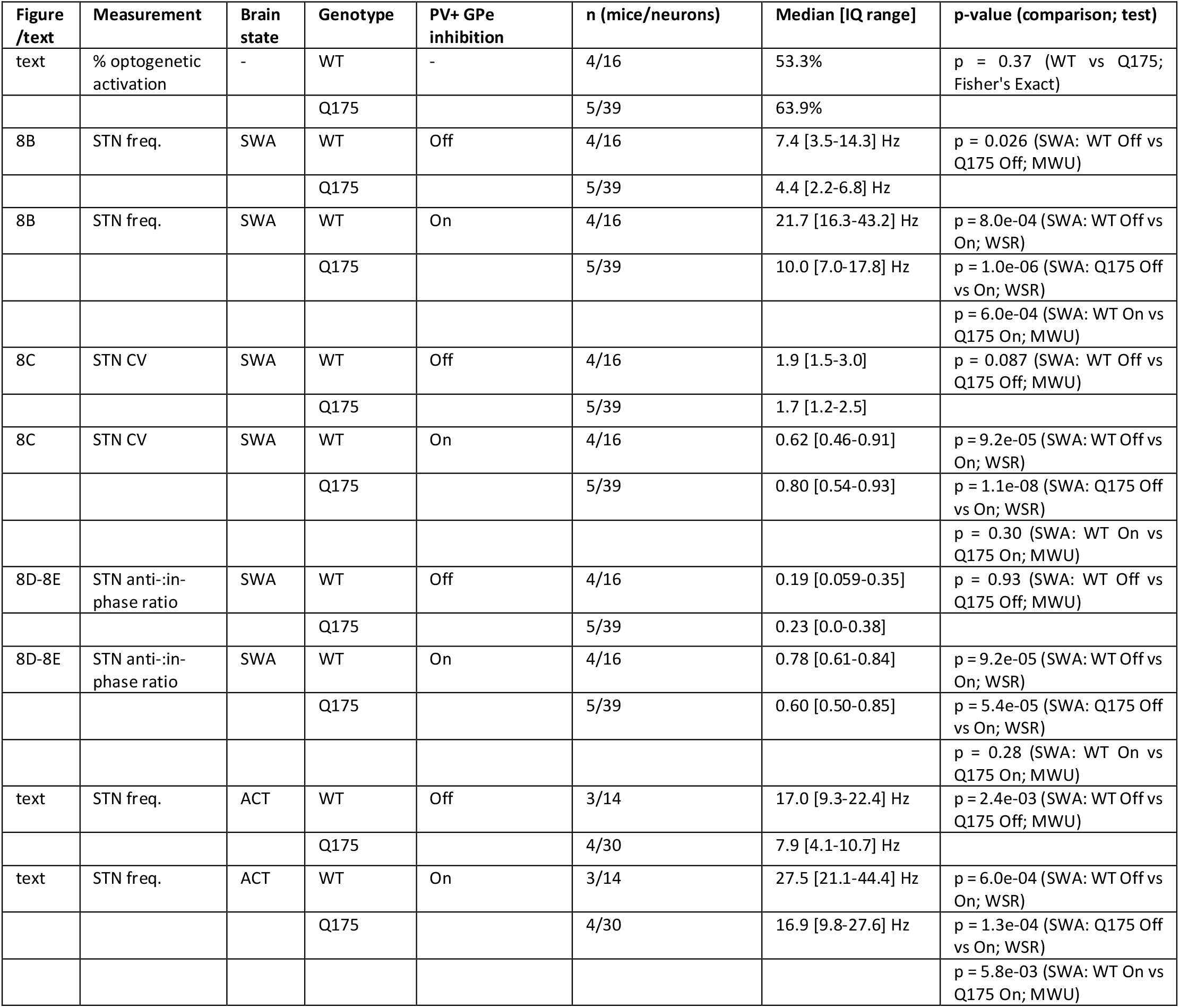

| Figure /text | Measurement | Brain state | Genotype | PV+ GPe inhibition | n (mice/neurons) | Median [IQ range] | p-value (comparison; test) |
| --- | --- | --- | --- | --- | --- | --- | --- |
| text | % optogenetic activation | - | WT | - | 4/16 | 53.3% | p = 0.37 (WT vs Q175; Fisher's Exact) |
|  |  |  | Q175 |  | 5/39 | 63.9% |  |
| 8B | STN freq. | SWA | WT | Off | 4/16 | 7.4 [3.5-14.3] Hz | p = 0.026 (SWA: WT Off vs Q175 Off; MWU) |
|  |  |  | Q175 |  | 5/39 | 4.4 [2.2-6.8] Hz |  |
| 8B | STN freq. | SWA | WT | On | 4/16 | 21.7 [16.3-43.2] Hz | p = 8.0e-04 (SWA: WT Off vs On; WSR) |
|  |  |  | Q175 |  | 5/39 | 10.0 [7.0-17.8] Hz | p = 1.0e-06 (SWA: Q175 Off vs On; WSR) |
|  |  |  |  |  |  |  | p = 6.0e-04 (SWA: WT On vs Q175 On; MWU) |
| 8C | STN CV | SWA | WT | Off | 4/16 | 1.9 [1.5-3.0] | p = 0.087 (SWA: WT Off vs Q175 Off; MWU) |
|  |  |  | Q175 |  | 5/39 | 1.7 [1.2-2.5] |  |
| 8C | STN CV | SWA | WT | On | 4/16 | 0.62 [0.46-0.91] | p = 9.2e-05 (SWA: WT Off vs On; WSR) |
|  |  |  | Q175 |  | 5/39 | 0.80 [0.54-0.93] | p = 1.1e-08 (SWA: Q175 Off vs On; WSR) |
|  |  |  |  |  |  |  | p = 0.30 (SWA: WT On vs Q175 On; MWU) |
| 8D-8E | STN anti-in-phase ratio | SWA | WT | Off | 4/16 | 0.19 [0.059-0.35] | p = 0.93 (SWA: WT Off vs Q175 Off; MWU) |
|  |  |  | Q175 |  | 5/39 | 0.23 [0.0-0.38] |  |
| 8D-8E | STN anti-in-phase ratio | SWA | WT | On | 4/16 | 0.78 [0.61-0.84] | p = 9.2e-05 (SWA: WT Off vs On; WSR) |
|  |  |  | Q175 |  | 5/39 | 0.60 [0.50-0.85] | p = 5.4e-05 (SWA: Q175 Off vs On; WSR) |
|  |  |  |  |  |  |  | p = 0.28 (SWA: WT On vs Q175 On; MWU) |
| text | STN freq. | ACT | WT | Off | 3/14 | 17.0 [9.3-22.4] Hz | p = 2.4e-03 (SWA: WT Off vs Q175 Off; MWU) |
|  |  |  | Q175 |  | 4/30 | 7.9 [4.1-10.7] Hz |  |
| text | STN freq. | ACT | WT | On | 3/14 | 27.5 [21.1-44.4] Hz | p = 6.0e-04 (SWA: WT Off vs On; WSR) |
|  |  |  | Q175 |  | 4/30 | 16.9 [9.8-27.6] Hz | p = 1.3e-04 (SWA: Q175 Off vs On; WSR) |
|  |  |  |  |  |  |  | p = 5.8e-03 (SWA: WT On vs Q175 On; MWU) |

**Figure 8.**
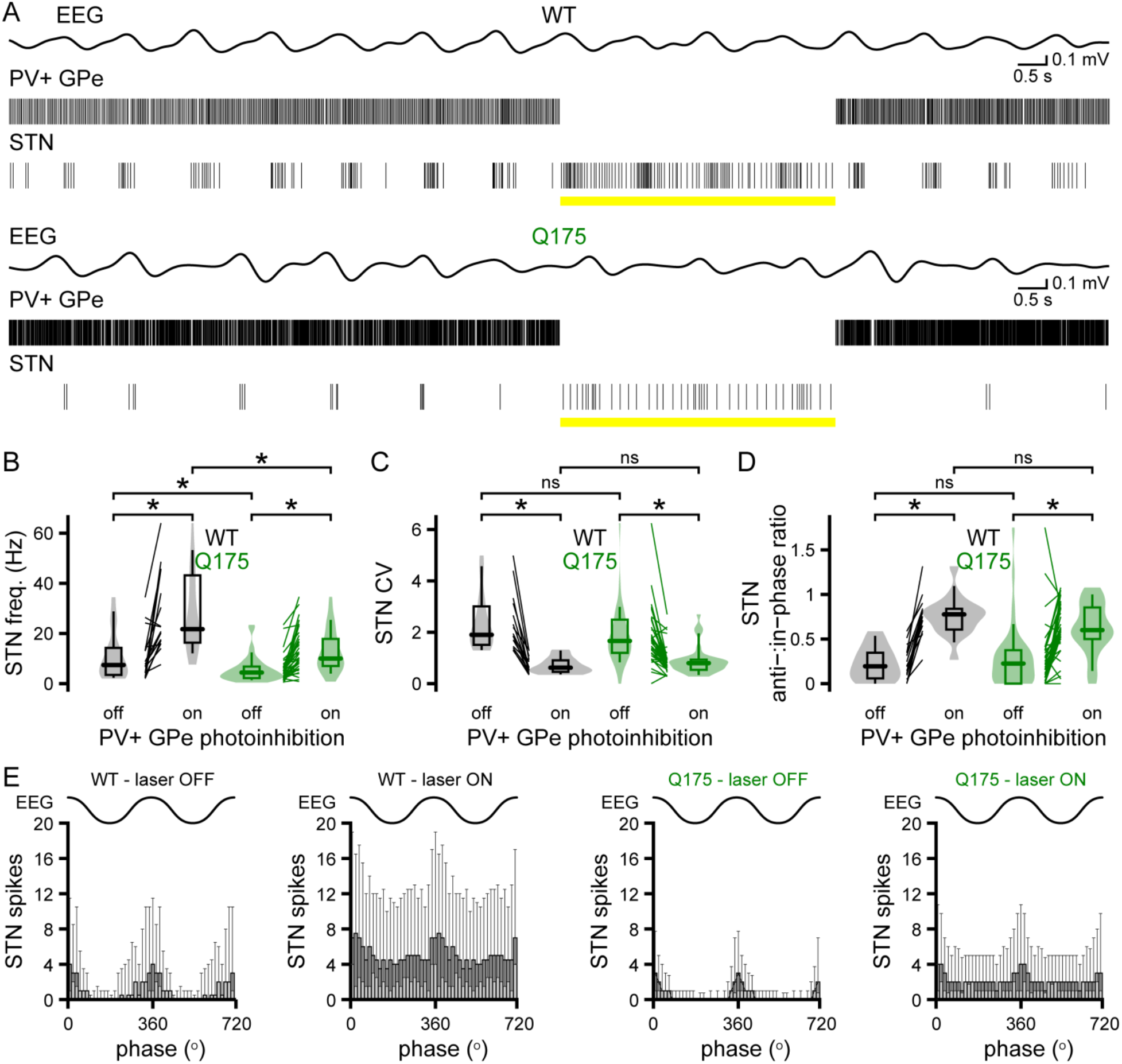
STN hypoactivity is partially rescued by optogenetic inhibition of PV+ GPe neurons in Q175 mice. (**A**) Representative examples of the EEG, band-pass filtered at 0.1-1.5 Hz, and concurrent PV+ GPe and responsive STN neuron activity in WT and Q175 mice before, during, and after optogenetic inhibition (yellow bar) of PV+ GPe neurons. (**A-C**) Optogenetic inhibition of PV+ GPe neurons disinhibited and regularized STN activity in both genotypes. However, the relative hypoactivity of STN neurons in Q175 mice was not fully reversed by optogenetic inhibition of PV+ GPe neurons (**B-C**, population data; WT, n = 16 neurons; Q175, n = 39 neurons). (**D-E**) The ratio of anti- to in-phase STN activity was increased to a similar extent by inhibition of PV+ GPe neurons in WT and Q175 mice (**D**, population data; **E**, population spike phase histograms). *, p < 0.05. ns, not significant.

## Discussion

Using cell class-specific optical identification and interrogation in conjunction with *in vivo* and *ex vivo* electrophysiology, we report widespread dysregulation of basal ganglia indirect pathway and arkypallidal neuron activity in Q175 mice at early symptomatic stages, 6-12 months prior to cell loss. Although, the EEG and layer V cortical projection neuron activity were similar in Q175 and WT mice, neuronal activity was markedly altered in the downstream basal ganglia. Relative to WT mice, D2-SPNs were hypoactive, prototypic PV+ GPe neurons were hyperactive, and putative arkypallidal and STN neurons were hypoactive in Q175 mice. Furthermore, PV+ GPe neurons exhibited upregulated autonomous firing *ex vivo*, and their hyperactivity was responsible, at least in part, for the hypoactivity of putative arkypallidal and STN neurons *in vivo* in Q175 mice.

### Cortical network and layer V projection neuron activity

The cortex of Q175 mice exhibits a variety of alterations, including nuclear aggregates of mHTT (Smith et al., 2014; Carty et al., 2015), reduced expression of TrkB receptors (Smith et al., 2014) and mGlu receptors (Bertoglio et al., 2018), and an increase in the ratio of inhibitory to excitatory transmission (Indersmitten et al., 2015). Cortical volume loss is also detectible at 4 months of age using MRI (Heikkinen et al., 2012) and at 12 months using stereological approaches (Smith et al., 2014). Despite these changes, we found that the EEGs of 6-month-old Q175 and WT mice were similar during both cortical SWA and ACT, suggesting that *in vivo* cortical network activity and depth of anesthesia in the two genotypes were comparable. We also discovered that the firing rate of identified layer V PT and putative IT cortical pyramidal neurons, many of which project to the basal ganglia (Harris and Shepherd, 2015), were similar in Q175 and WT mice during both cortical SWA and ACT. Consistent with our observations, the firing rate of unidentified cortical neurons is also not altered in awake full-length and other knock-in HD mouse models (Rebec, 2018). The firing patterns of identified layer V PT and putative IT cortical pyramidal neurons during SWA and ACT were also similar in Q175 and WT mice. However, in awake behaving HD mice, burst firing and correlated activity are diminished in unidentified cortical neurons during both rest and movement (Rebec, 2018). The differences between our and other studies likely reflects different network dynamics in anesthetized versus awake mice and measurements from identified versus unidentified cortical neurons.

### Striatal projection neuron activity

Consistent with direct synaptic driving by cortical pyramidal neurons, optogenetically identified D2-SPNs and putative D1-SPNs discharged predominantly during the active phase of the slow cortical oscillation in both Q175 and WT mice. During cortical SWA, the median frequency of D2-SPN activity in Q175 mice was only half of that in WT mice. However, during sensory-evoked cortical desynchronization, D2-SPN activity was almost completely suppressed in both Q175 and WT mice. Although putative D1-SPN activity was similar in Q175 and WT mice during cortical SWA, it was relatively elevated in Q175 mice during ACT. Overall, putative D1-SPN activity was elevated relative to D2-SPN activity during cortical SWA in Q175 mice only. No differences in the precision of D2-SPN and putative D1-SPN activity were observed in Q175 and WT mice. Given the similar rate of cortical projection neuron activity in WT and Q175 mice during cortical SWA, the relative hypoactivity of D2-SPNs in Q175 mice is consistent with previously reported reductions in cortico-striatal axospinous synapses, cortico-striatal synaptic LTP, cortico-striatal synaptic strength, and distal dendritic excitability in D2-SPNs (Plotkin and Surmeier, 2015; Sebastianutto et al., 2017; Carrillo-Reid et al., 2019) (but see Goodliffe et al., 2018). In contrast, the relative retention of cortico-striatal synaptic transmission and plasticity, and integrative properties in D1-SPNs (Plotkin and Surmeier, 2015; Carrillo-Reid et al., 2019) presumably contributed to their relatively normal/elevated activity in Q175 mice. While, the depolarization of the resting membrane potential and increased axosomatic excitability of SPNs in Q175 mice (Heikkinen et al., 2012; Indersmitten et al., 2015; Beaumont et al., 2016; Sebastianutto et al., 2017; Goodliffe et al., 2018) appear not to compensate for reduced cortical excitation of D2-SPNs, they may contribute to the augmentation of D1-SPN activity during cortical ACT. Consistent with impaired cortical driving of D2-SPNs *in vivo*, firing in response to cortical stimulation, burst firing, and correlated activity are diminished in unidentified striatal neurons in HD mice (Beaumont et al., 2016; Rebec, 2018). The spontaneous activity of unidentified striatal neurons has also been reported to be elevated, reduced, or unchanged in anesthetized knock-in or awake full-length HD mice (Miller et al., 2008; Estrada-Sanchez et al., 2015; Beaumont et al., 2016). Differences between these studies and our data presumably reflect the use or absence of cell-identification approaches and/or recordings in anesthetized or awake mice. The responses of SPNs to cortical excitation are also heavily sculpted by GABAergic striatal interneurons through both tonic and feedforward inhibition (Tepper et al., 2004; Tepper et al., 2018). Thus, recent *ex vivo* studies (Holley et al., 2019a; Holley et al., 2019b), which reveal alterations in interneuron excitability and their inhibition of SPNs argue that striatal interneuron dysfunction may also contribute to the abnormal activity of SPNs in HD mice.

### Prototypic PV+ GPe and arkypallidal neuron activity

During cortical SWA prototypic PV+ GPe neurons and PV-GPe, putative arkypallidal neurons were hyperactive and hypoactive, respectively, in Q175 mice. The hyperactivity of prototypic PV+ GPe neurons *in vivo* could reflect the hypoactivity of upstream D2-SPNs, which are relatively huge in number (Oorschot, 1996). However, the reduction in the frequency of striatopallidal transmission in HD mice may also be partially compensated by an increase in transmission strength, as reported recently (Perez-Rosello et al., 2019). Indeed, during cortical SWA, PV+ GPe neurons in Q175 mice often exhibited reduced firing during D2-SPN activity. It is not clear whether an increase in vesicular GABA content and/or increased expression of postsynaptic receptors underlies the increase in D2-SPN to prototypic PV+ GPe neuron transmission strength. However, increased expression of GPe GABA_A_ receptors has been widely reported in HD (Glass et al., 2000; Waldvogel et al., 2015). Prototypic PV+ GPe neurons were also clearly hyperactive when upstream D2-SPN activity was negligible, as evinced by their increased antiphasic activity. Thus, the hyperactivity of prototypic PV+ GPe neurons *in vivo* may stem in part from their elevated autonomous firing, which we observed in *ex vivo* brain slices derived from Q175 mice. Given the widespread nature and potency of prototypic PV+ GPe neuron projections throughout the basal ganglia (Bevan et al., 1998; Mallet et al., 2012; Mastro et al., 2014; Abdi et al., 2015), it will be critical to determine which ion channel mechanisms are responsible for their upregulated autonomous firing in Q175 mice, and whether upregulated autonomous activity results from the cell-autonomous effects of mutant huntingtin or reflects homeostatic compensation, e.g. in response to STN hypoactivity.

The hypoactivity of putative arkypallidal neurons in Q175 mice appears to be caused by excessive inhibition emanating from hyperactive prototypic PV+ GPe neurons because arkypallidal hypoactivity was fully alleviated by optogenetic inhibition of prototypic PV+ GPe neurons *in vivo*. The regulation of “arkypallidal” neuron activity by PV+ GPe neurons *in vivo* is consistent with powerful GABAergic inhibition of arkypallidal neurons by prototypic GPe neurons (Mallet et al., 2012; Aristieta et al., 2020; Ketzef and Silberberg, 2020). However, optogenetic silencing of prototypic PV+ GPe neurons also disinhibited hypoactive STN neurons, and given that STN neurons innervate arkypallidal neurons (Aristieta et al., 2020; Pamukcu et al., 2020), it is also possible that STN disinhibition contributed to the rescue of arkypallidal neuron firing in Q175 mice. The finding that the autonomous firing of putative arkypallidal neurons was unaffected in brain slices from Q175 mice further suggests that synaptic mechanisms are responsible for the hypoactivity of arkypallidal neurons in Q175 mice. The low rate of putative arkypallidal neuron autonomous firing in *ex vivo* brain slices from both Q175 and WT mice compared to identified prototypic PV+ GPe neurons is consistent with previous reports in WT mice (Abdi et al., 2015; Hernandez et al., 2015). Whether prototypic PV+ GPe or STN neuron to arkypallidal neuron transmission is normal, upregulated, or downregulated in Q175 mice remains to be determined. Although, the majority of prototypic GPe neurons express PV and the vast majority of arkypallidal neurons express FoxP2, a minority of GPe neurons express neither marker (Mastro et al., 2014; Abdi et al., 2015; Dodson et al., 2015; Hernandez et al., 2015). Thus, a subset of the putative arkypallidal neurons that we recorded from could include PV-prototypic GPe neurons and/or GPe neurons with non-classical projection patterns (Mastro et al., 2014; Abecassis et al., 2020). Finally, our data contrast with another study that reported no change in the frequency of firing in GPe neurons in anesthetized Q175 mice (Beaumont et al., 2016). In that study the identity of the GPe neurons that were recorded and the relative depth of anesthesia were not known, potentially occluding the alterations reported here.

### Subthalamic nucleus activity

In Q175 mice we found that STN activity was less than half of that in WT mice during both cortical SWA and ACT. The hypoactivity of STN neurons observed here is analogous to that reported in Q175 mice in which the EEG was not measured and the cortical activity state was therefore unknown (Beaumont et al., 2016), and more robust than activity changes in YAC128 (Callahan and Abercrombie, 2015b) and R6/2 mice (Callahan and Abercrombie, 2015a), in which only a subset of STN neurons exhibited diminished activity. The relative hypoactivity of STN neurons in Q175 mice could arise from a variety of sources, including hyperactivity of upstream prototypic PV+ GPe neurons, loss of autonomous STN activity (Atherton et al., 2016), and reduced cortical excitation (Beaumont et al., 2016). Indeed, STN hypoactivity was significantly but only partially alleviated in Q175 mice during optogenetic inhibition of PV+ GPe neurons, consistent with excessive inhibition from upstream prototypic GPe neurons, but also diminished autonomous STN activity and possibly cortical driving. During cortical SWA, STN activity is relatively phase-locked to the active component, arguing that it is cortically generated. Indeed, physical removal of cortical input or depression of cortical activity eliminates this pattern of STN activity in WT rodents (Magill et al., 2001). Interestingly phase-locking of STN activity to cortical SWA was unaffected in Q175 mice both prior to and following optogenetic inhibition of prototypic GPe neurons. Together these observations argue that the combination of excessive inhibition arising from prototypic GPe neurons and impaired autonomous firing are largely responsible for the abnormal frequency and pattern of STN activity in Q175 mice. That said, how the synaptic properties of GPe-STN and cortico-STN inputs are affected in Q175 mice is currently unknown.

As suggested above, the hypoactivity of STN neurons may contribute to the hypoactivity of arkypallidal neurons in Q175 mice because optogenetic inhibition of presynaptic prototypic GPe neurons and disinhibition of the STN rescued arkypallidal neuron activity. In contrast, STN hypoactivity did not prevent hyperactivity of upstream prototypic GPe neurons, presumably because the hyperactivity of prototypic GPe neurons resulted from a combination of elevated autonomous firing and reduced transmission from hypoactive D2-SPNs.

### Functional implications

According to the classic rate model of basal ganglia encoding, hypoactivity of D2-SPNs, and arkypallidal and STN neurons, and hyperactivity of prototypic GPe neurons in Q175 mice should elevate movement (Albin et al., 1989; Kravitz et al., 2010). Indeed, the firing rate changes that we report here are largely opposite to those reported in anesthetized and awake Parkinson’s disease rats and mice (Magill et al., 2001; Mallet et al., 2008a; Mallet et al., 2008b; Sharott et al., 2017; Ryan et al., 2018; Kovaleski et al., 2020). However, 6-month-old Q175 mice exhibit akinesia and bradykinesia that is similar to but less severe than in Parkinsonism (Heikkinen et al., 2012; Menalled et al., 2012; Pouladi et al., 2013; Kosior and Leavitt, 2018). Given the large size of their trinucleotide repeat expansion, Q175 mice may therefore model joHD in which bradykinesia and akinesia present in the absence of the initial hyperkinetic/dyskinetic phase seen in aoHD (Fusilli et al., 2018; Tereshchenko et al., 2019).

Recent studies also argue that the indirect pathway is not solely devoted to movement suppression. Co-activity of D2- and D1-SPNs is necessary for normal volitional movement (Cui et al., 2013; Tecuapetla et al., 2016; Markowitz et al., 2018; Parker et al., 2018). Furthermore, increases in arkypallidal and STN neuron activity, possibly due to D2-SPN inhibition of prototypic GPe neurons, have been variably linked to the execution, inhibition, and prevention of movement (Dodson et al., 2015; Mallet et al., 2016; Pasquereau and Turner, 2017; Aristieta et al., 2020). Thus, dysregulation of the indirect pathway may actively contribute to the motor impairments seen in Q175 mice. Indeed, our unpublished data indicate that movement-associated increases in STN activity are profoundly diminished in awake, behaving Q175 mice.

A final outstanding question is whether 6-month-old Q175 mice better mimic prodromal HD (Paulsen, 2010; Long et al., 2014; Reilmann et al., 2014) because the abnormal indirect pathway activity reported here precedes the cell loss that is traditionally thought to underlie the disease’s clinical presentation (Reiner et al., 1988; Albin et al., 1992; Richfield et al., 1995; Sapp et al., 1995; Vonsattel and DiFiglia, 1998). If 6-month-old Q175 mice do indeed model prodromal HD, our data argue that indirect pathway neurons are profoundly dysregulated in the presymptomatic stage of the disease, long before they are lost.

## Materials and methods

### Animals

Procedures were performed in compliance with the policies of the National Institutes of Health and approved by the Institutional Animal Care and Use Committee of Northwestern University. Mice were maintained on a 14-hour light/10-hour dark cycle with food and water ad libitum, and monitored regularly by animal care technicians, veterinarians, and research staff.

Heterozygous Q175 mice (B6J.129S1-*Htt*^*tm1*.*1Mfc*^/190ChdiJ; RRID: IMSR_JAX: 029928) were bred with homozygous PT-kj18-cre mice (Tg(Sim1-cre)KJ18Gsat; RRID: MMRRC_031742-UCD), or A2A-cre mice (Tg(Adora2a-cre)KG139Gsat; RRID: MMRRC_036158-UCD), or PV-cre mice (B6.Cg-*Pvalb*^*tm1*.*1(cre)Aibs*^/J; RRID: IMSR_JAX: 012358) to generate offspring that were either homozygous for HTT (WT), or heterozygous for HTT and mHTT (Q175), and heterozygous for cre-recombinase. The following experimental mice were used (median and age range are reported): WT/PT-kj18-cre: age = 233, 225-239 days old, n = 5; Q175/PT-kj18-cre: age = 238, 228-244 days old, n = 4; WT/A2A-cre: age = 210, 210-233 days old, n = 3; Q175/A2A-cre: age = 211, 211-235 days old, n = 3; WT/PV-cre: age = 219.5, 204-232 days old, n = 4; Q175/PV-cre: age = 220, 205-234 days old, n = 5). Experimental mice were male except for PV-cre mice, where both male (WT/PV-cre: n = 3; Q175/PV-cre: n = 3) and female (WT/PV-cre: n = 1; Q175/PV-cre: n = 2) mice were used. Data from male and female PV-cre mice were overlapping and therefore pooled.

### Stereotaxic injection of viral vectors

Anesthesia was induced with vaporized 3-4% isoflurane (Smiths Medical ASD, Inc., Dublin, OH, USA) followed by an intraperitoneal (IP) injection of ketamine (100 mg/kg). After securing the mouse in a stereotaxic instrument (Neurostar, Tubingen, Germany), anesthesia was maintained with 1-2% isoflurane. AAVs diluted in HEPES buffered synthetic interstitial fluid (HBS SIF: 140 mM NaCl, 23 mM glucose, 15 mM HEPES, 3 mM KCl, 1.5 mM MgCl_2_, 1.6 mM CaCl_2_; pH 7.2 with NaOH; 300–310 mOsm/L) were then injected under stereotaxic guidance. AAV injection was conducted over 5-10 minutes at each site. An additional 5-10 minutes was then allowed for the injectate to diffuse prior to syringe retraction. ChR2(H134R)-eYFP was virally expressed in: 1) layer V PT neurons in PT-kj18-cre mice through unilateral injection of AAV9.EF1a.DIO.hChR2(H134R)-eYFP.WPRE.hGH (RRID: Addgene_20298; 1 × 10^13^ genome copies/mL; AP: +0.6 mm, +1.2 mm, +1.8 mm; ML: 1.5 mm; DV: 1.0 mm; 0.5 μl per injection) 2) D2-SPNs in A2A-cre mice through unilateral injections of AAV9.EF1a.DIO.hChR2(H134R)-eYFP.WPRE.hGH (RRID: Addgene_20298; 3 × 10^11^ genome copies/mL; AP: +0.4 mm, +0.9 mm; ML: 2.2 mm; DV: 3.7 mm, 2.7 mm; 0.3 μl per injection). Arch-GFP was virally expressed in PV+ GPe neurons in PV-cre mice through unilateral injection of AAV9.CBA.Flex.Arch-GFP.WPRE.SV40 (RRID: Addgene_22222; 5 × 10^11^ genome copies/mL; AP: −0.27 mm; ML: 1.90 mm; DV: 3.95 mm, 3.45 mm; 0.25 μl per ventral injection and 0.20 μl per dorsal injection). eGFP was virally expressed in PV+ GPe neurons in PV-cre mice through unilateral injection of AAV9.Syn.DIO.eGFP.WPRE.hGH (RRID: Addgene_100043; 3 × 10^11^ genome copies/mL; AP: −0.27 mm; ML: 1.90 mm; DV: 3.95 mm, 3.45 mm; 0.25 μl per ventral injection and 0.20 μl per dorsal injection).

### In vivo electrophysiological recording

2-4 weeks following surgery anesthesia was induced with vaporized 3-4% isoflurane followed by IP injection of urethane (1.5 g/kg, IP; Sigma-Aldrich, St. Louis, MO, USA). Mice were placed back into their home cage for approximately 60 minutes until the toe-pinch withdrawal reflex was abolished. If necessary additional urethane supplements (0.5 g/kg, IP) were administered every 30 minutes until the withdrawal reflex was eliminated. Mice were then placed into a stereotaxic instrument (David Kopf Instruments, Tujunga, CA, USA) for the duration of the recording session with urethane supplements administered as required to maintain anesthesia. Craniotomies were drilled over the primary motor cortex (AP: +1.4 mm; ML: 1.5 mm), striatum (AP: +0.65 mm; ML: 1.95 mm), GPe (AP: −0.3 mm; ML: 2.0 mm), and/or STN (AP: − 1.9 mm; ML: 1.4 mm) and irrigated with HBS. The intracranial EEG was recorded from a peridural screw electrode (MS-51960-1; McMaster-Carr, Chicago, IL, USA) affixed over the ipsilateral primary motor cortex of A2A-cre and PV-cre mice. In PT-kj18-cre mice in which extracellular recordings of motor cortical neurons were made, the EEG screw was implanted over the contralateral primary motor cortex. Extracellular single unit recordings were acquired using silicon tetrodes/optrodes (A1×4-tet-10mm-100-121-A16 and A1×4-tet-10mm-100-121-OA16, respectively; NeuroNexus Technologies, Ann Arbor, MI, USA) connected to a 64-channel Digital Lynx (Neuralynx, Bozeman, MT, USA) data acquisition system via a unity gain headstage, with a reference wire implanted adjacent to the ipsilateral temporal musculature. Signals were sampled at 40 kHz, with a gain of 14 X. Online digital FIR filters were applied. Single unit activity was band-pass filtered between 200-9000 Hz and EEG and LFP signals were band-pass filtered between 0.1-400 Hz. Optogenetic stimulation was delivered using either a 473 nm diode laser (LuxX+ 473-100; Omicron-Laserage Laserprodukte GmbH, Rodgau, Germany) or a custom 577 nm laser system (Genesis MX STM 577-500 OPSL CW; Coherent Inc., Santa Clara, CA, USA). In order to histologically verify recording sites, silicon tetrodes/optrodes were dipped in a lipophilic florescent dye (DiI; 20 mg/ml in 50% acetone/methanol; D282; ThermoFisher Scientific, Waltham, MA, USA) prior to implantation. Sensory-evoked cortical ACT was generated by pinching the hindpaw for 5 seconds using a pair of fine forceps (Fine Science Tools, Foster City, CA, USA). ChR2(H134R)-expressing neurons were optogenetically stimulated through exposure to 473 nm light (< 6 mW) for a duration of 5 ms. Stimulation was repeated at least 5 times with each trial of stimulation separated by a minimum of 2 minutes. Arch-GFP-expressing neurons were optogenetically inhibited through delivery of 577 nm light (< 6 mW) for a duration of 5 seconds. Optogenetic inhibition was repeated at least 2 times with each trial being separated by a minimum of 2 minutes. Laser intensity was calibrated as power at the tip of the optrode prior to probe implantation and was verified at the conclusion of each experiment.

### Immunohistochemistry

To enable reconstruction of electrode tracks and confirm specificity of viral expression, mice were given a lethal dose of anesthetic and then perfused transcardially with ∼5-10 ml of 0.01 M phosphate-buffered saline (PBS, pH 7.4; P3813, MilliporeSigma, Darmstadt, Germany) followed by 15-30 ml of 4% paraformaldehyde (PFA) in 0.1 M phosphate buffer (PB, pH 7.4). Each brain was then carefully removed and post-fixed overnight in 4% PFA (prepared in 0.1 M PB, pH 7.4) before being washed in PBS, blocked, and sectioned in the coronal plane at 70 µm using a vibratome (VT1000S; Leica Microsystems Inc., Richmond, Illinois, USA). Sections were then processed for the immunohistochemical detection of NeuN, an antigen expressed by neurons that is commonly used to delineate brain structures. Thus, sections were washed in PBS and incubated for 48-72 hr at 4°C in anti-NeuN (1:200; clone A60; MilliporeSigma; RRID: AB_2532109) in PBS with 0.5% Triton X-100 (MilliporeSigma) and 2% normal donkey serum (Jackson ImmunoResearch, West Grove, PA, USA). Sections were then washed in PBS and incubated for 90 min at room temperature in Alexa Fluor 488- or 594-conjugated donkey anti-mouse IgG (1:250; Jackson ImmunoResearch; RRID: AB_2313584; RRID: AB_2340621) in PBS with 0.5% Triton X-100 and 2% normal donkey serum. In a subset of PV-cre mice, in which PV+ GPe neurons expressed Arch-GFP or eGFP, adjacent sections of the GPe were processed for the immunohistochemical detection of PV (primary antibody: 1:1000 guinea pig anti-PV; Synaptic Systems, Gottingen, Germany; RRID: AB_2156476; secondary antibody: 1:250 Alexa Fluor 594 donkey anti-guinea pig IgG; Jackson ImmunoResearch; RRID: AB_2340474) or FoxP2 (primary antibody: 1:1000 rabbit anti-FoxP2; Millipore Sigma; RRID: AB_1078909; secondary antibody: 1:250 Alexa Fluor 594 donkey anti-rabbit IgG; Jackson ImmunoResearch; RRID: AB_2340621). Sections were washed in PBS and then mounted on glass slides with ProLong Diamond Antifade Reagent (P36965; ThermoFisher Scientific, Waltham, MA, USA). Mountant was allowed to cure for at least 24 hours prior to storage at 4°C or imaging. DiI and immunofluorescent labelling were imaged using a Zeiss Axioskop 2 microscope (Zeiss, Oberkochen, Germany), equipped with an Axiocam CCD camera (426508-9901-000: Zeiss), and Neurolucida software (MFB Bioscience, Williston, VT, USA). Finally, representative images were acquired using confocal laser scanning microscopy (A1R; Nikon Instruments Inc., Melville, NY, USA).

### In vivo electrophysiological analysis

Estimates of spectral power density were extracted using the Chronux data analysis toolbox (Bokil et al., 2010) for MATLAB (chronux.org; MathWorks, Natick, MA, USA). The EEG signal was down-sampled to 1000 Hz and spectral power was assessed at a resolution frequency of 0.061 Hz. Putative single unit activity was discriminated with Plexon Offline Sorter software (Version 3; Plexon. Inc., Dallas, TX, USA; RRID: SCR_000012) using a combination of template matching, principal component analysis, and manual clustering. To ensure putative single unit isolation, spike timestamps were aligned to the peak of the action potential and a threshold of < 0.5% ISI within 2 ms with a dead time of 500 µs was required (% ISI within 2 ms for all experiments; cortical neurons: WT: 0.0, 0.0-0.0, n = 22; Q175: 0.0, 0.0-0.0, n = 36; striatal neurons: WT: 0.0, 0.0-0.0, n = 83; Q175: 0.0, 0.0-0.0, n = 61; GPe neurons: WT: 0.08, 0.02-0.29, n = 50; Q175: 0.05, 0.01-0.24, n = 55; STN neurons: WT: 0.23, 0.11-0.38, n = 30; Q175: 0.11, 0.00-0.19, n = 61; values represent median and interquartile range). Electrophysiological data were visually inspected in NeuroExplorer (Nex Technologies, Colorado Springs, CO; RRID: SCR_001818) and then exported to MATLAB (MathWorks, Natick, MA; RRID: SCR_001622). Epochs with stable and robust SWA or ACT were selected for analysis. Neurons were identified as D2-SPNs if an action potential was elicited within 10 ms of ChR2(H134R) stimulation in at least 3 of 5 trials. The direct and downstream impact of optogenetic inhibition of PV+ GPe neurons was assessed from neuronal activity prior to and during 577 nm illumination. Neurons were deemed responsive to optogenetic inhibition if the firing rate or CV of PV+ GPe neurons or their postsynaptic targets were consistently altered by at least 2 standard deviations of mean pre-stimulus values.

To ensure that measurements were made from areas with opsin expression, analysis was restricted to recordings in which there was at least one responsive neuron on a given tetrode array. Mean firing rates were calculated from the number of spikes divided by epoch length. The coefficient of variation of the interspike interval (CV) was used as a metric of regularity. To examine the relationship between cortical SWA and neuronal firing, phase histograms were generated in MATLAB. The EEG signal was down-sampled to 1000 Hz and SWA was extracted by applying a bandpass 0.1-1.5 Hz 2nd order Butterworth filter in the forward and reverse directions (to avoid phase shifts). The instantaneous phase of the EEG was calculated from the Hilbert transform (Le Van Quyen et al., 2001). In order to correct for the non-sinusoidal nature of slow cortical oscillations, the empirical cumulative distribution function (MATLAB) was applied (Siapas et al., 2005; Mallet et al., 2008b; Abdi et al., 2015; Kovaleski et al., 2020). Thus, each spike was assigned to a phase of the EEG from 0-360° (with 0°/360° and 180° corresponding to the peak active and inactive components of the EEG, respectively). Data were binned at 15° and anti-:in-phase ratio measurements were calculated. Population phase histograms are plotted as median and interquartile range.

### Ex vivo electrophysiological recording

Mice were lightly anesthetized with isoflurane, deeply anesthetized with ketamine/xylazine (87/13 mg/kg, IP), and then perfused transcardially with ∼10 ml of ice-cold sucrose-based artificial cerebrospinal fluid (sACSF; 230 mM sucrose, 2.5 mM KCl, 1.25 mM NaH_2_PO_4_, 0.5 mM CaCl_2_, 10 mM MgSO_4_, 10 mM glucose, 26 mM NaHCO_3,_ 1 mM sodium pyruvate, 5 µM L-glutathione; equilibrated with 95% O_2_ and 5% CO_2_). The brain was then removed, immersed in ice-cold 95% O_2_/5% CO_2_-equilibrated sACSF and sectioned at 250 µm in the sagittal plane with a vibratome (VT1200S; Leica Microsystems Inc., IL, USA). Slices were then transferred to a holding chamber, immersed in ACSF (126 mM NaCl, 2.5 mM KCl, 1.25 mM NaH_2_PO_4_, 2 mM CaCl_2_, 2 mM MgSO_4_, 10 mM glucose, 26 mM NaHCO_3_, 1 mM sodium pyruvate, 5 µM L-glutathione; equilibrated with 95% O_2_ and 5% CO_2_), held at 35°C for 30 min, and then maintained at room temperature. Individual brain slices were transferred to a recording chamber where they were perfused at 4–5 ml/min with synthetic interstitial fluid (SIF; 126 mM NaCl, 3 mM KCl, 1.25 mM NaH_2_PO_4_, 1.6 mM CaCl_2_, 1.5 mM MgSO_4_, 10 mM glucose, 26 mM NaHCO_3_; equilibrated with 95% O_2_ and 5% CO_2_) at 35°C. Patch clamp recordings were made using 3-6 MΩ impedance, borosilicate glass electrodes filled with HBS SIF. Electrodes were positioned under visual guidance (Axioskop FS2, Carl Zeiss) using computer-controlled micromanipulators (Luigs and Neumann, Ratingen, Germany). Somatic recordings were made in the loose-seal, cell-attached configuration using an amplifier (MultiClamp 700B; Molecular Devices, Sunnyvale, CA, USA), and associated digitizer (Digidata 1440A; Molecular Devices) controlled by pCLAMP 10.3 (Molecular Devices). Electrode capacitance was compensated and signals were low-pass filtered online at 10 kHz and sampled at 25 kHz. Recordings of autonomous action potential generation were made in the presence of 20 µM DNQX, 50 µM D-AP5, 10 µM Gabazine, and 2 µM CGP 55845 to block synaptic transmission at AMPA, NMDA, GABA_A_, and GABA_B_ receptors, respectively. All drugs used were purchased from Hello Bio Inc. (Princeton, NJ, USA) and bath applied. PV+ GPe neurons were identified through visualization of eGFP under 460 nm LED epifluorescent illumination (OptoLED; Cairn Research, Faversham, Kent, UK). PV-GPe neurons were also recorded if eGFP-positive PV+ GPe neurons were in the same field of view to ensure that recording sites were made from areas with reporter expression. The frequency and regularity of GPe neuron activity were calculated from 30 second recording epochs.

## Acknowledgements

This study was funded by CHDI and NIH NINDS grant R37 NS041280. Confocal imaging work was performed at the Northwestern University Center for Advanced Microscopy, which is supported by NIH NCI grant CCSG P30 CA060553 (awarded to the Robert H. Lurie Comprehensive Cancer Center). The authors thank S. Ulrich, D.R. Schowalter, and B.M. Erjavec for the maintenance and supply of transgenic mice for this study.

## Declaration of Interests

The authors declare no competing interests.

## Notes

### Competing Interest Statement

The authors have declared no competing interest.

